# The *Myb* family genes in the rice pathogen *Magnaporthe oryzae*: Finding and deleting more family members involved in pathogenicity

**DOI:** 10.1101/2021.12.28.474317

**Authors:** Ya Li, Xiuxia Zheng, Mengtian Pei, Mengting Chen, Shengnan Zhang, Chenyu Liang, Luyao Gao, Pin Huang, Stefan Olsson

**Author notes:** Correspondence: Ya Li and Stefan Olsson, &.

## Abstract

Proteins with DNA binding Myb domains have been suggested in regulating development and stress responses. *Magnaporthe oryzae* is considered the most destructive pathogen of rice. We screened the genome for genes with Myb domains encoding since these can be needed for pathogenesis. We found *Myb1-19*. Only MoMyb1 was previously shown to be involved in pathogenesis. We succeeded in deleting 12 of the other 18 genes. MoMyb2 deletion affected mainly growth, while MoMyb13 or MoMyb15 deletions gave additional defects in conidiation and plant infection. However, RT-qPCR showed that none of the 19 Myb genes are negligibly expressed. Instead, they have different expression profiles hours post-infection when infecting rice plants. Considering this, the unchanged infection phenotype for 9 gene deletions surprised us, and we extended the analysis to expression co-regulation of all 19 Myb proteins and found 5 co-regulated groups of predicted Myb-domain proteins. MoMyb13 or MoMyb15 are discussed and motivated as candidates for further, more detailed studies with aims also outside of plant pathology. Referring to what is found in other eukaryotes, we finally discuss possible redundancy or compensatory regulations for many of the other Myb genes hiding or compensating for the effect of many complete deletions.

**IMPORTANCE:** *Magnaporthe oryzae* is considered the most important rice pathogen limiting rice production. Our study attempts to find all genes encoding a DNA-binding gene family called Myb, and we found 19, many of which have not been studied before. The Myb gene family is suspected to regulate stress responses the pathogen needs to overcome plant defenses. Inhibiting or disturbing these genes, if they are indeed regulatory, can open new ways of controlling the pathogen and learning more about its physiology and ecology.

## INTRODUCTION

Rice blast disease, caused by *Magnaporthe oryzae*, is one of the most destructive diseases on rice worldwide. Each year, this disease leads to estimated economic damage of $66 billion, which is enough to feed 60 million people (Pennisi, 2010). The disease initiates from the *M. oryzae* conidia contacting to rice surface, then the conidia germinate and develop a structure called appressoria for host penetration. The mature appressoria accumulate high internal turgor pressure (Howard et al., 1991) and mechanically penetrate rice cells by forming a hyphal penetration peg (Kankanala et al., 2007). Once completing colonization, multiple infection hyphae expand intra- and intercellularly in rice tissue, and the typical blast lesions appear in 3 to 5 days (Sakulkoo et al., 2018). New conidia are produced from lesions and released to start new infection cycles. Besides attacking rice, *M. oryzae* can infect wheat, barley, finger millet, and foxtail millet (Gladieux et al., 2018). Recently, this fungus caused a wheat blast outbreak in Bangladesh, resulting in huge losses (Islam et al., 2016; Malaker et al., 2016). Given the economic importance, genetic tractability, and genome sequence availability, *M. oryzae* has become a model to study the fungal pathogenesis and interaction with host plants (Dean et al., 2005; Ebbole, 2007). Understanding the transcription-factor-mediated cellular or biological processes of *M. oryzae* is beneficial to develop novel and practical strategies to control the blast disease and ensure global food security.

Based on the similarity of the DNA-binding domain, transcription factors are categorized into up to 61 families, including bZIP, bHLH, C2H2 zinc finger, homeobox, Zn2Cys6, Myb, etc. (Verma et al., 2017). The Myb domain family of proteins is one of the largest and most diverse families containing transcription factors. A highly conserved Myb DNA-binding domain characterizes these proteins (Ambawat et al., 2013; Du et al., 2009; Prouse and Campbell, 2012; Roy, 2016; Verma et al., 2017; L., Wang et al., 2018; X., Wang et al., 2018). The Myb domain generally consists of 1 to 3 imperfect amino acid repeats; each repeat contains 50 to 53 amino acids and spatially constitutes a helix turn helix structure. The name Myb domain was acquired from v-Myb, the oncogenic motif of avian myeloblastosis virus (AMV), where it was first discovered (X., Wang et al., 2018). The Myb family is present in all eukaryotes and could thus have more than 1 billion years of evolutionary history (Eme et al., 2017). In plants, the Myb family proteins are mainly considered as transcription factors or repressors (Allan and Espley, 2018; Ambawat et al., 2013; Baldoni et al., 2015; Cao et al., 2020; Dong et al., 2015; Du et al., 2009; Dubos et al., 2010; Kim et al., 2014; Li et al., 2019; Li et al., 2019; Lin et al., 2011; Lin et al., 2012; Liu et al., 2015; Ma and Constabel, 2019; Mateos et al., 2006; Matheis et al., 2017; Mitra, 2018; Pattabiraman and Gonda, 2013; Prouse and Campbell, 2012; Roy, 2016; Valsecchi et al., 2017; Verma et al., 2017; X., Wang, et al., 2018; Wieser and Adams, 1995). However, proteins that are not classical transcription factors can contain Myb-type DNA binding domains (Hogues et al., 2008; Ma and Constabel, 2019; Mukherjee et al., 2012; Pattabiraman and Gonda, 2013; Ramos-Sáenz et al., 2019; Rodríguez-Sánchez et al., 2011; Roy, 2016; X., Wang et al., 2018; Yu et al., 2010). The Myb family does not have many members comprising 4 to 5 proteins in animal genomes (Prouse and Campbell, 2012). While in plants, the Myb family has expanded to 100 to 200 members (Prouse and Campbell, 2012). The fungal Myb family size is smaller than in plants but more extensive than in animals, with 10 to 50 members per genome (Verma et al., 2017). Animal Myb proteins are reported to regulate cell division and a discrete subset of cellular differentiation events (X., Wang et al., 2018). Plant Myb proteins bind DNA and regulate many metabolic, cellular, and developmental processes as transcription factors or in other ways (Prouse and Campbell, 2012; Roy, 2016). Fungal DNA-binding proteins of the Myb family have been implicated to be essential for regulation to withstand stresses (Verma et al., 2017; L., Wang et al., 2018). They should consequently play a role in the pathogenicity of plant pathogens (Verma et al., 2017), especially during the entry to, and in, the necrotrophic phase of fungal plant pathogens starting as biotrophs, like *M. oryzae* when the plant starts to detect the intruding fungus and actively counteract it by innate immune responses producing ROS and toxic metabolites (Cao et al., 2022). In *M. oryzae*, 13 Myb family genes have previously been identified (Verma et al., 2017). We searched for more to investigate if any of them in in addition to Myb1 and Myb8 (Dong et al., 2015; Lee et al., 2021) singly have critical roles in regulating the pathogenicity of *M. oryzae* on rice since Myb proteins seems essential for managing fungal stress (Verma et al., 2017; L., Wang et al., 2018) characteristic of plant infection.

First, we searched for the Myb-like domains in annotated *M. oryzae*-proteins. We found 19 genes encoding for proteins containing Myb DNA binding domains, including *Myb1*, previously proven to be involved in pathogenicity (Dong et al., 2015). We attempted to delete all the remaining 18 genes except Myb1 and succeeded in deleting 12 genes. Only 2 of the 12 genes we deleted strongly affected pathogenicity when deleted singly, although all 19 identified Myb domain genes were relatively highly expressed *in planta* during different stages of infection. One of the 2 genes affecting pathogenicity is regulating melanin biosynthesis. Therefore, the lack of standard panel phenotypic effect of the deletions of many of the Myb-domain genes highly expressed during plant infection is discussed.

## RESULTS

### 1. Myb family domain protein identification in *M. oryzae* and analysis of their amino acid similarity

Genes for 19 *Myb* family DNA binding proteins were predicted in *M. oryzae* by the Fungal Transcription Factor Database (FTFD, http://ftfd.snu.ac.kr/index.php?a=view). One gene was previously reported as MoMyb1 (Dong et al., 2015), and 9 more recently identified as *MoMyb2-10* (Lee et al., 2021) while we were writing up this work. The other 9 we named *MoMyb11* to *19* in this study (**Fig. 1A** and **B**). Protein size and domain structure analysis showed that the 19 MoMyb protein sizes vary from the maximum 2305 aa to the minimum 250 aa. Each protein consists of at least 1 Myb domain and up to 3 (**Fig. 1A**). According to amino acid sequence similarities, the 19 *M. oryzae* Myb domain-containing proteins cluster into four main groups that could have different functions (**Fig. 1B**). To investigate some of the possible biological functions of MoMyb-proteins and especially if some of the Myb proteins, other than MoMyb1 and MoMyb8 (Dong et al., 2015; Lee et al., 2021), that has been shown to be involved in the pathogenesis of rice in *M. oryzae*, 12 Myb gene deletion mutants (*Δmomyb2, 3, 4, 5, 6, 8, 9, 10, 13, 15, 16, and 18*) were generated and confirmed by Southern blot (**Fig S1**). Mutants of 5 other *Myb* genes (*MoMyb7, 11, 12, 17, 19*) could not be obtained, even when screened from more than 200 transformants of each gene containing the gene fragment that potentially can replace the target gene. *MoMyb7* has been deleted before (Lee et al., 2021) but is not involved in pathogeneses. Even if we could not achieve a mutation for this one, our lack of mutations indicates that *MoMyb17, 19, 11, and 12* could be essential genes.

**Figure 1.**
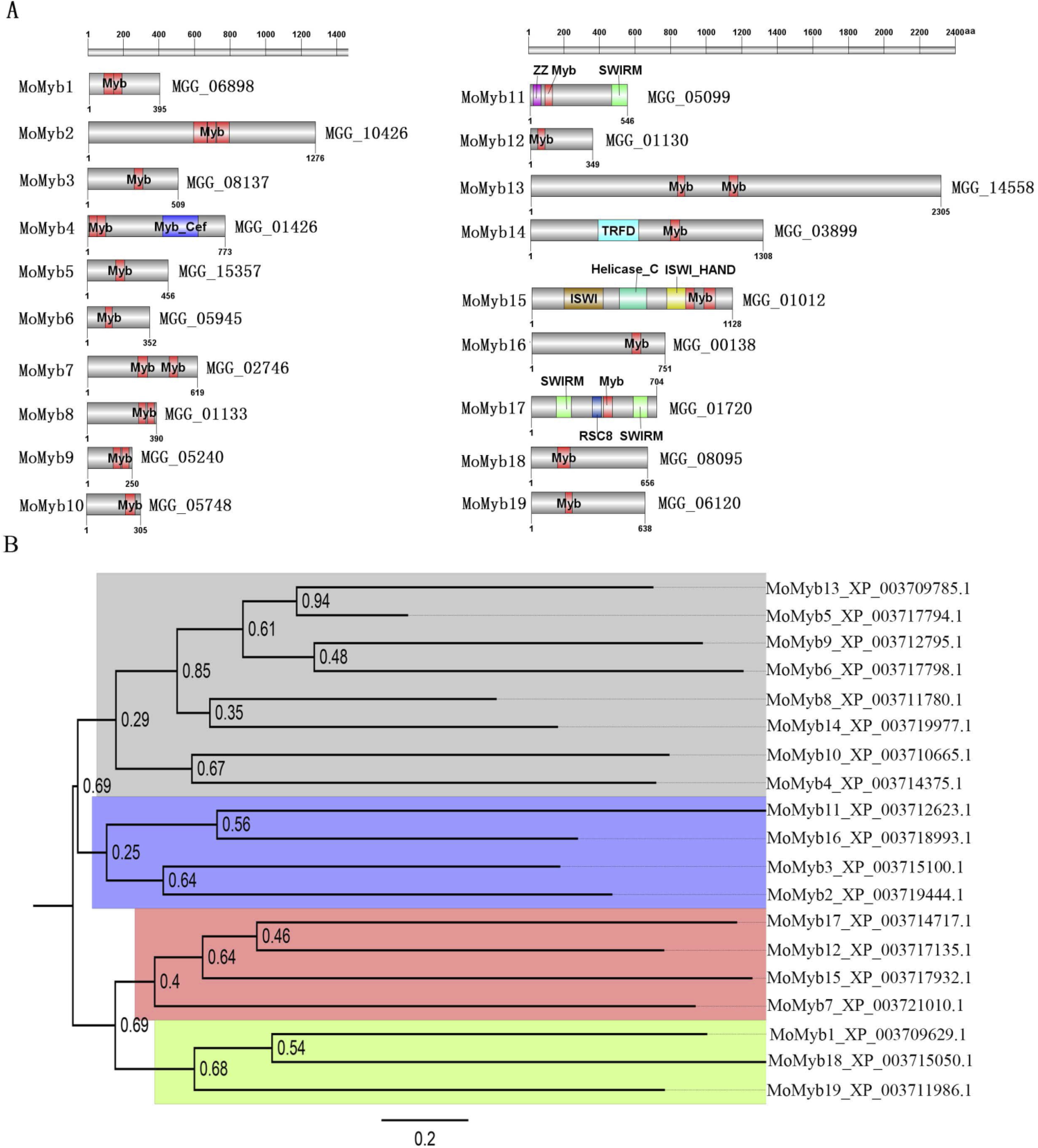
Domain structure of the identified genes encoding for Myb domain-containing proteins and their sequence similarities. (**A**) Identified conserved domains in the found Myb domain-containing proteins. (**B**) The similarity analysis of predicted amino acid sequences showed that the 19 proteins could be sorted into 4 groups marked by the color code.

### 2. MoMyb2, 13, and 15 are involved in regulating *M. oryzae* growth

A growth assay was performed by growing the mutants on three types of media, including complete medium (CM), minimal medium (MM), and rice bran medium (RBM), for ten days to test whether the respective Myb-proteins, affect *M. oryzae* radial growth. We found 3 MoMyb mutants lacking proteins MoMyb2, 13, and 15 (*Δmomyb2-101, Δmomyb13-11*, and *Δmomyb15-7*) that showed significantly decreased colony size on all three media as compared to the background isolate Ku80 and the corresponding gene complementary strains (*Δmomyb2-101/MoMyb2, Δmomyb13-11/MoMyb13*, and *Δmomyb15-7/MoMyb15*) (**Fig. 2A and B**). In contrast, the other 9 Myb mutants showed the same colony size as the Ku80 strain (**Fig. S2**), suggesting that MoMyb2, 13, and 15 directly or indirectly regulate some aspects of *M. oryzae* growth. The growth inhibition rate of these three mutants on different media was also calculated. The result showed that the mutants show different growth inhibition effects on different media. *Δmomyb13-11* showed the highest inhibition on RBM, followed by MM and CK; *Δmomyb2-101* showed the highest inhibition on CM, followed by MM and RBM; *Δmomyb15-7* showed the highest inhibition on MM, followed by CM and RBM (**Fig. 2C**). These results suggest that the different nutrients of these media exerted different stresses on these 3 Myb mutants.

**Figure 2.**
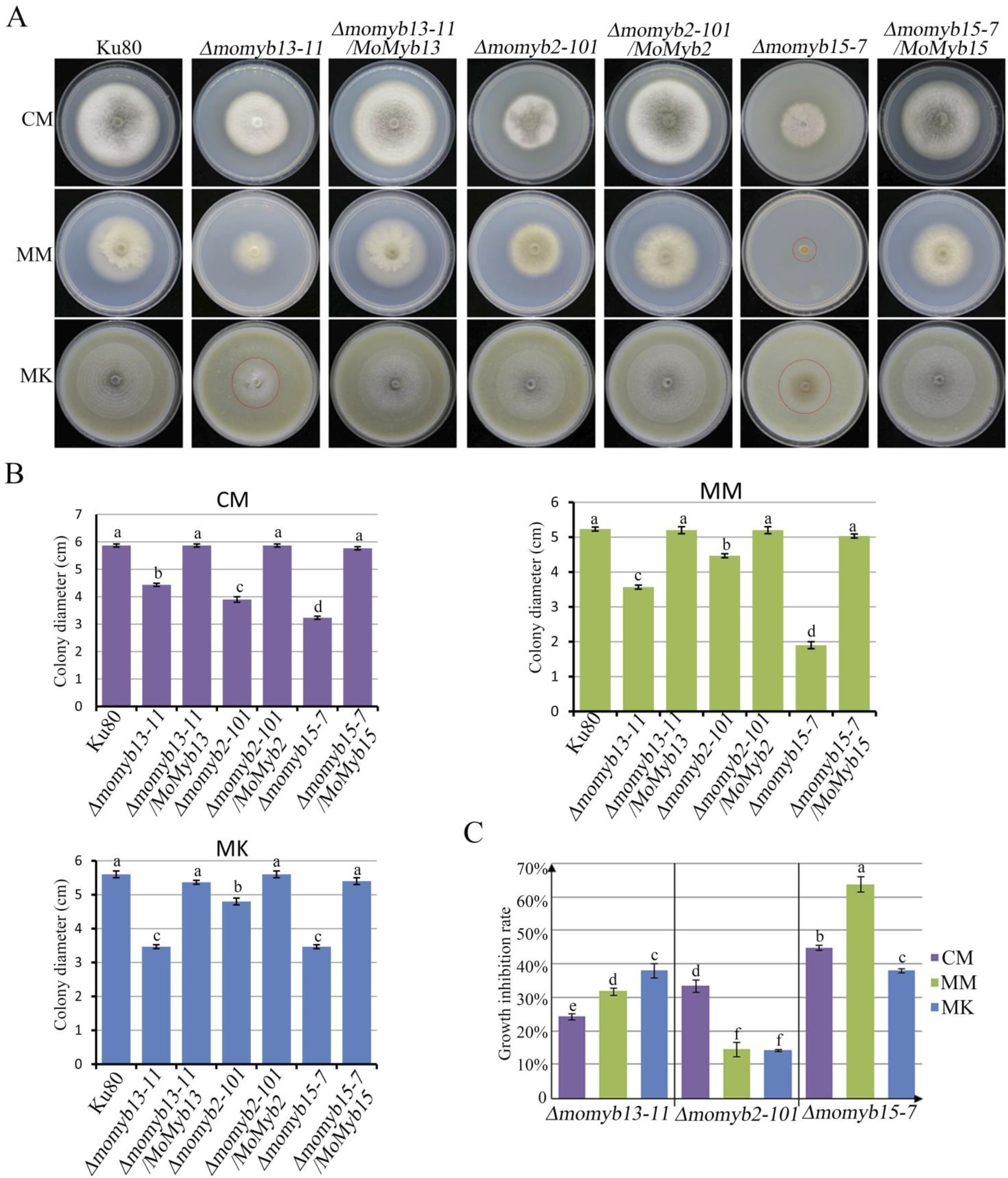
The radial growth rate of *Δmomyb2, Δmomyb13*, and *Δmomyb15* are slower than in the background strain Ku80. (**A**) Images of the morphology of the different strains on the three different media, complete medium (CM), minimal medium (MM), and rice bran medium (RBM). (**B**) Measured colony diameter after 10day incubation in alternating dark/light (12h/12h) at 25 °C. (**C**) Growth inhibition in deletion strains was measured as colony diameter after 10-day incubation in alternating dark/light (12h/12h) at 25 °C compared to the background strain Ku80. Bars indicate SEM and bars with the same letters are not significantly different (P>0.05; t-test).

### 3. MoMyb13 and 15 appear to be involved in *M. oryzae* pathogenicity in rice

We tested the 12 *Myb* gene mutations we had achieved for pathogenicity on rice leaves using inoculation with mycelia on agar blocks since none of the *Δmomyb13* and *Δmomyb15* that affected growth formed conidia (see below). Only *Δmomyb13* and Δmomyb15 had reduced pathogenicity when agar block inoculation was performed on both intact or wounded barley and rice leaves, and complementations with their respective gene restored pathogenicity (**Fig. 3A-D**). Conidia inoculation could not be used for these two mutants since no conidia were formed, as seen in a conidiation assay (**Fig. 4A and B**). The lack of pathogenicity does not appear to result from a lack of appressoria formed by the mycelial inoculum since appressoria were formed (**Fig. 3E**). There were no effects on pathogenicity for the other mutants of the remaining 10 *MoMyb* genes we managed to delete. Those had all typical conidia that could be used for the pathogenicity assay (**Fig. S3**).

**Figure 3.**
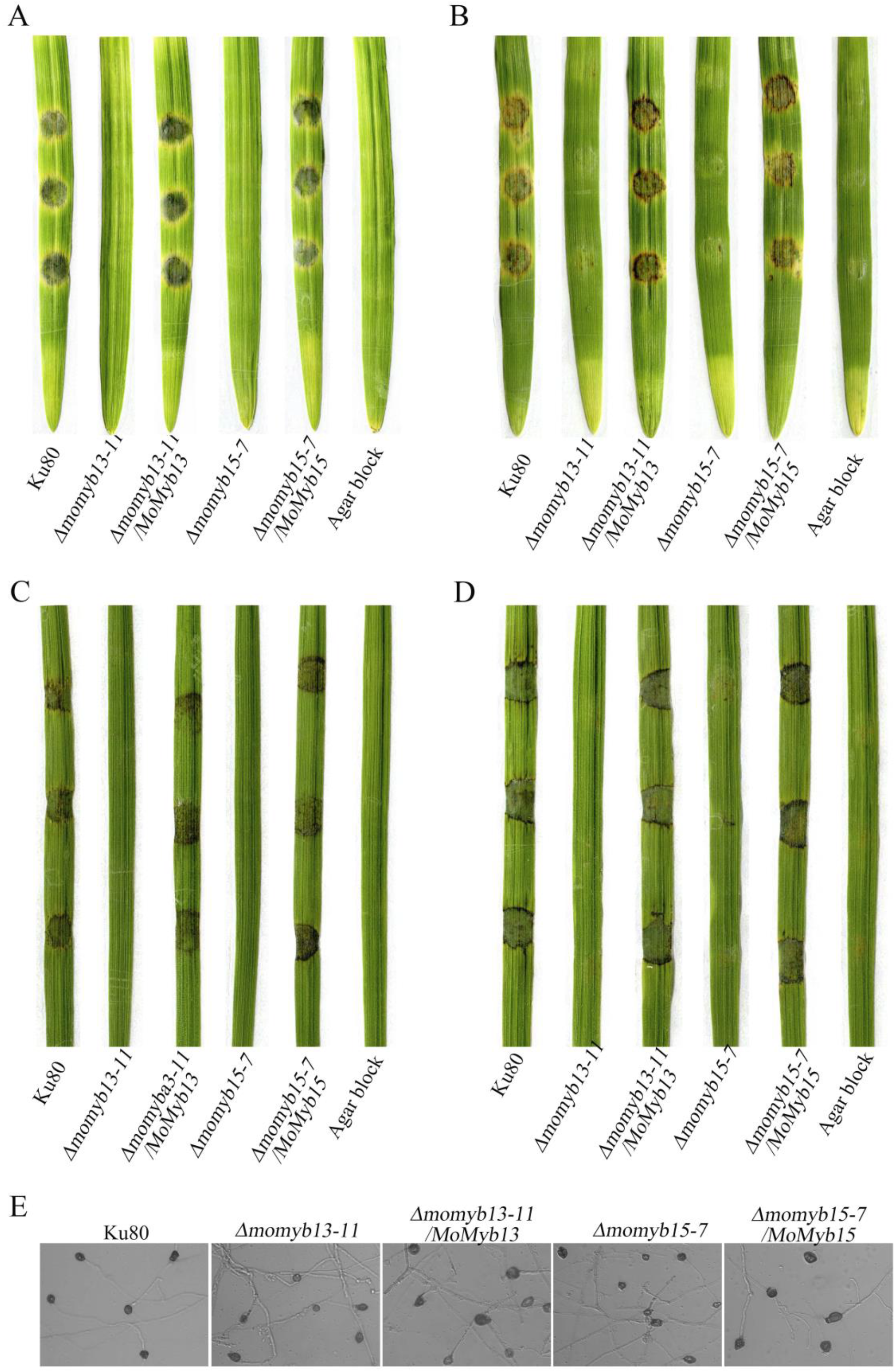
MoMyb13 and MoMyb15 are involved in *M. oryzae* pathogenicity since mutants of the genes showed reduced lesions when tested with the agar block technique on excised barley and rice leaves, but this lack of lesion is not the result of lack of mycelia appressorium formation. Test of mutants and background strain on intact (**A**) and wounded (**B**) barley leaves. Test of mutants and background strain on intact (**C**) and wounded (**D**) rice leaves. Mycelia appressorium formation from mycelial fragments on a hydrophobic surface (**E**).

**Figure 4.**
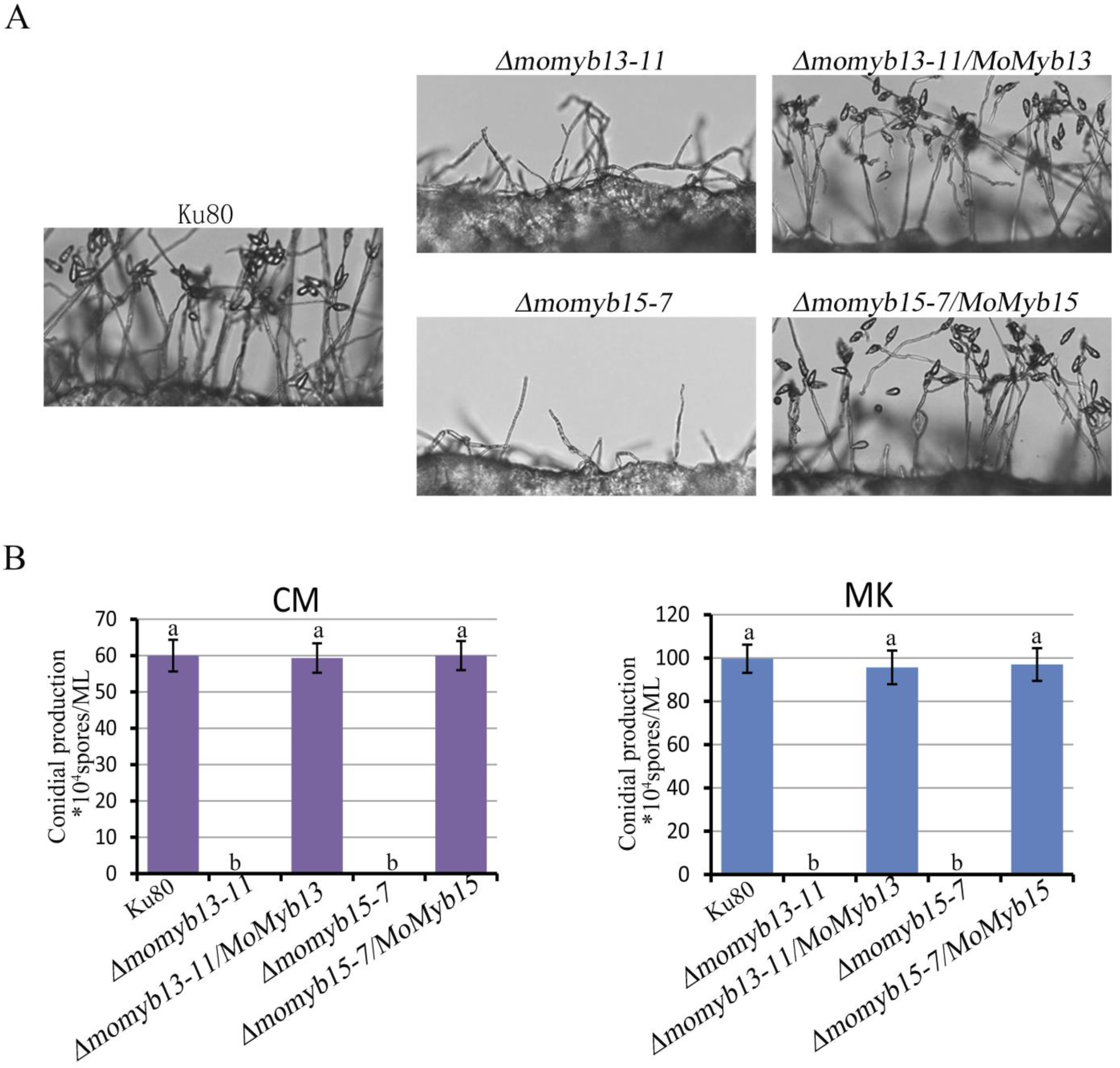
*Δmomyb13* and *Δmomyb15* make no conidia, while Ku80 and complement strains readily make. (**A**) Conidiophore morphology and conidiation. (**B**) Conidia production on CM and RBM.

### 4. MoMyb13 and MoMyb15 are involved in conidiation, and MoMyb13 regulates hydrophobins

Since conidia formation is essential for the fungus to spread rice blast disease from plant to plant, conidiation was more closely investigated. A conidia induction assay was performed on CM and RBM media to analyze the function of Myb proteins in *M. oryzae* conidiation. Conidia was also now absent for both mutants, and their respective complementation completely restored conidiation showing that both MoMyb13 and MoMyb15 are indeed necessary for conidiation, at least on these traditional growth media (**Fig. 4A** and **B**). Small hydrophobic proteins cover aerial mycelia of many ascomycetes, so-called hydrophobins, making aerial hyphae, including conidiophores, hydrophobic (Bayry et al., 2012; Berger and Sallada, 2019). We had noticed that the *Δmomyb13* had a more wettable mycelium (**Fig. 5A**) indicative of a lack of hydrophobins and thus decided to test if any described hydrophobin genes are downregulated in the *Δmomyb13* strain as these are not affected by direct regulation by MoMyb1 even if wettability was affected (Lee et al., 2021). Hydrophobin *MPG1* involved in pathogenesis (Beckerman, 1996) and affect wettability was downregulated in *Δmomyb13* (**Fig. 5B**). Artificial upregulation of *MPG1* in *Δmomyb13* (**Fig. 5C**) could not restore the original pathogenicity (**Fig. 5D**), indicating that MoMyb13 regulates more than hydrophobins necessary for successful conidia production and infection of rice.

**Figure 5.**
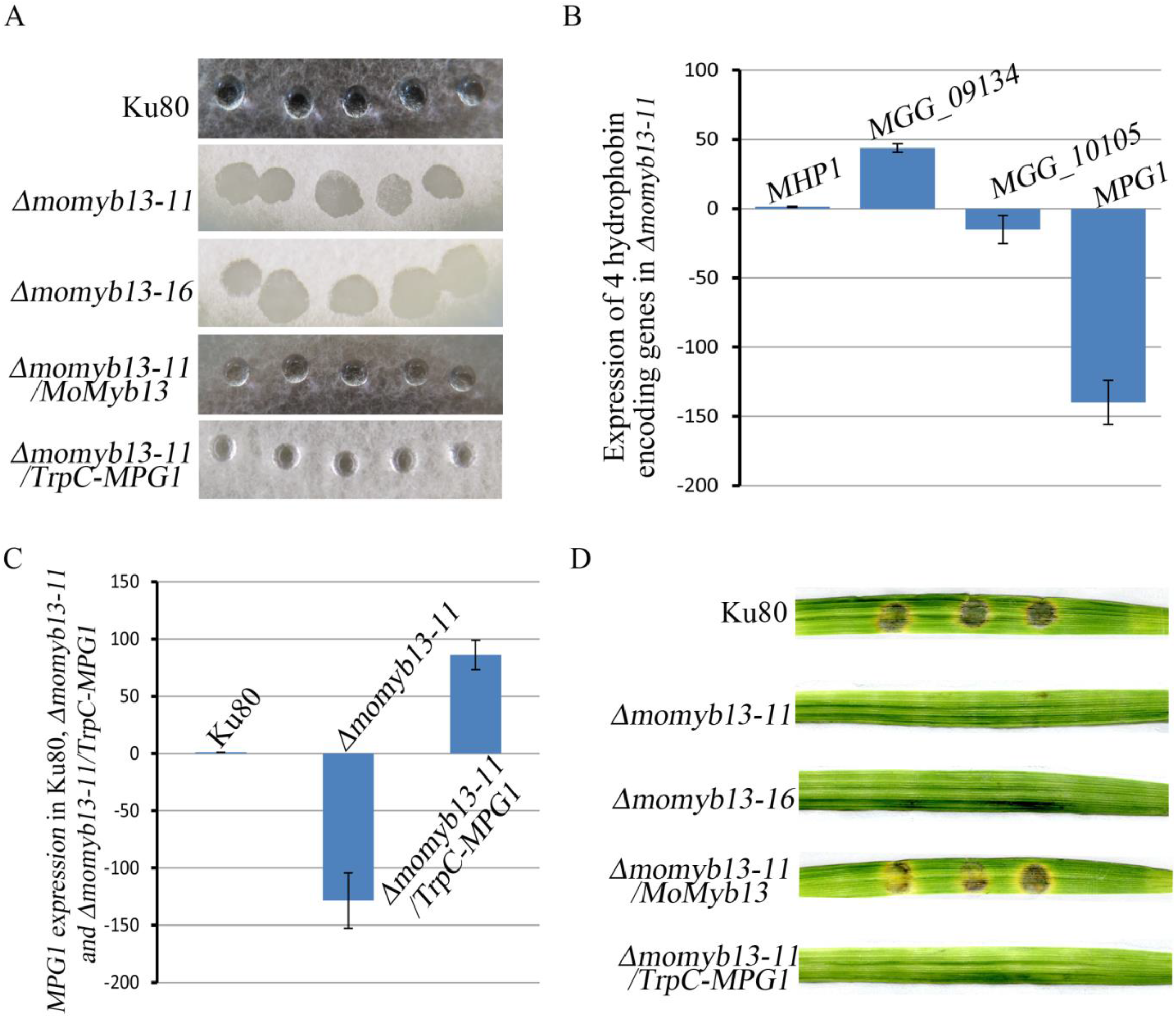
Hydrophobin production, necessary for producing hydrophobic arial conidia-producing mycelium, by MPG1 is downregulated in *Δmomyb13*, but overexpression of the MPG1 gene in the *Δmomyb13* strain does not restore pathogenicity. (**A**) colonies of *Δmomyb13* are easily wettable, while the surface of the background strain (Ku80), the complementation strain, or the *Δmomyb13* strain containing the overexpressed hydrophobin *MPG1 (Δmomyb13-11/TrpC-MPG1*) is not wettable. (**B**) Percent change in expression of 4 hydrophobin genes in *Δmomyb13* h compared to *Ku80*, showing that *MPG1* is severely downregulated while the others are less affected. (**C**) Percent change in *MPG1* expression in *Ku80, Δmomyb13 and Δmomyb13/TrpC-MPG1* strains compared to *Ku80*. (**D**) Leaf lesions formed by the *Ku80, Δmomyb13*, and *Δmomyb13/TrpC-MPG1* strains using the agar block technique. Error bars in (A) and (C) show the 95% confidence intervals. Bars with confidence intervals not overlapping with other bars’ confidence intervals or levels (as zero) are significantly different (P<0.05 for the null hypothesis that they are the same).

### 5. Sensitivity of Momyb13, Momyb2, and Momyb15 mutants to different stresses

A standard panel of stress treatments tested whether the MoMyb mutants affected growth rates on different stressing media (Li et al., 2014). A complete medium (CM) without additions was used as the control medium. The MoMyb13 mutant was inhibited by SDA, strongly inhibited by CR and H_2_O_2_ but stimulated by NaCl and SOR **(Fig. 6A and B)**. That can result from a weakened cell wall with less melanization (melanin is an antioxidant) combined with lower membrane permeability and higher activity of efflux pumps as compensation for these cell wall weaknesses in the mutant. MoMyb15 mutants are very inhibited by SDS affecting membrane integrity but also by NaCl **(Fig. 6A and B)** which suggest that membrane pumps depending on the membrane potential might not be necessarily upregulated and working in the mutant (Gostinčar et al., 2011). Loss of Myb2 mainly resulted in lower inhibition than the WT especially for CR suggesting that MoMyb2 is less important as stress regulator while MoMyb5 only showed increased inhibition for the SDS treatment leading to membrane disruptions. The results suggest that both MoMyb13 and MoMyb15 directly or indirectly regulate genes for proteins that strengthen the cell wall and cell membrane barrier in several ways important for growth inside a plant and might explain why deletion of these two genes lowered the pathogenicity of the fungus.

**Figure 6.**
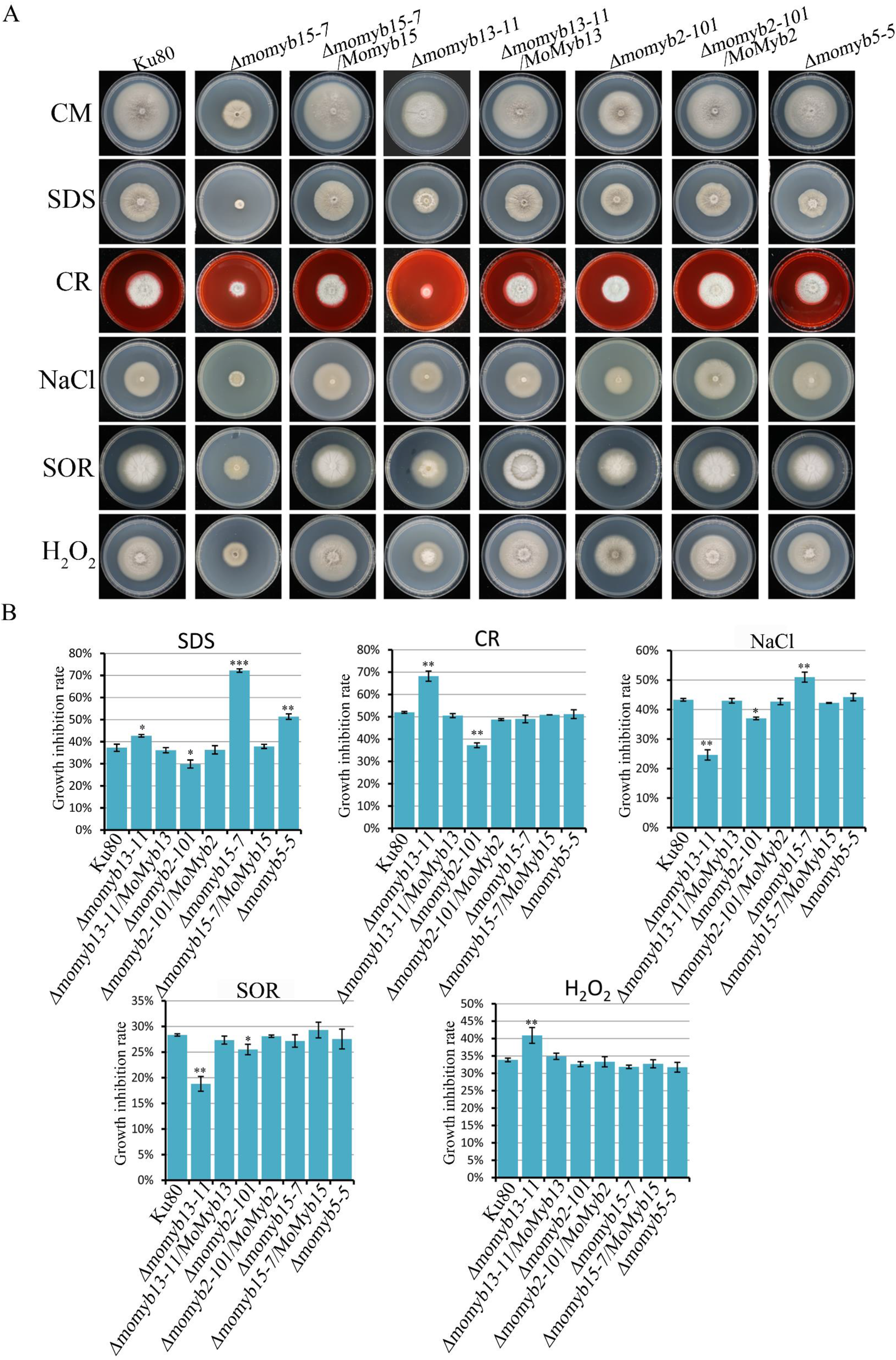
The three mutants, *Δmomyb5, Δmomyb13*, and *Δmomyb15*, showed reduced stress tolerance compared to the background strain Ku80 using a standard panel of stress treatments. (**A**) Colony size and morphology of Ku80, the mutants, and the complements after 10 days in alternating dark/light (12h/12h) at 25°C growing on CM or CM with additions of SDS, CR, NaCl SOR, or H2O2 as stresses (see methods). (**B**) Same as in (**A**) but shows the average of 3 replicates with error bars. Error bars show the standard error means of the respective averages, and stars mark statistical differences compared to Ku80.

### 6. MoMyb-GFP accumulates in the nucleus

All the 18 newly identified Myb-protein-encoding genes were expressed as *GFP*-containing fusion constructs, and the localization of these MoMyb protein GFP-fusions was investigated. It was confirmed that for all putative Myb protein fusions, we got a strong enough GFP signal to visualize through confocal microscopy; they accumulate to nucleus-like structures (**Fig. 7A-C** and **S4**). The three genes that showed at least some changed deletion phenotypes were investigated in more detail (**Fig. 7A-C**) in a Histone1-RFP background to label the nucleus in a different color (Zhang et al., 2019). It then became evident that the relatively small green “dots” of the MoMyb2-GFP indicate a localization to the nucleolus, where no other of the MoMybs seems to accumulate (**Fig. 7A-C** and **S4**).

**Figure 7.**
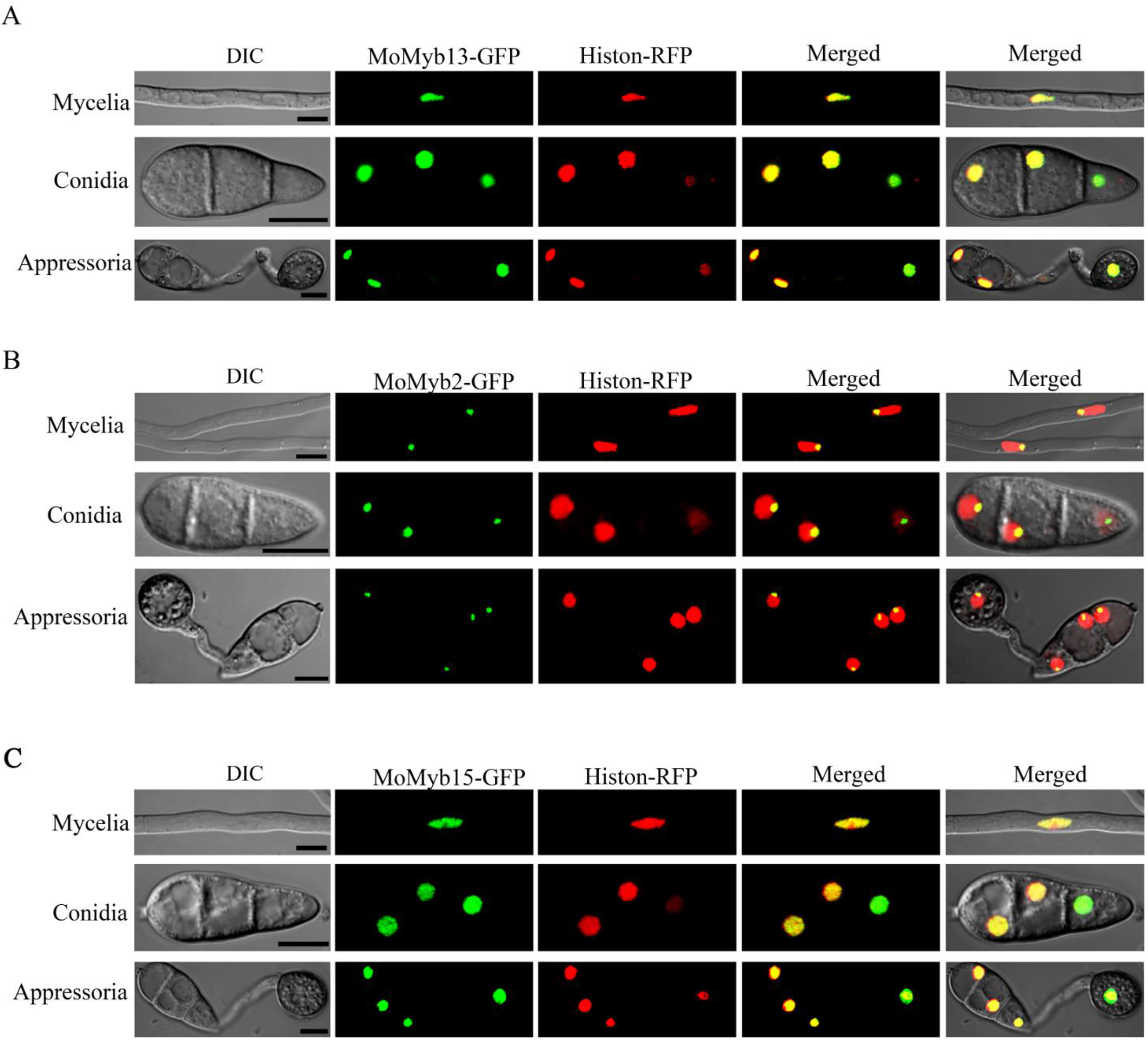
All three *momyb*-deletions that showed phenotypes differing from Ku80 (*Δmomyb13, Δmomyb2*, and *Δmomyb15*) localize to conidia, appressorial, and mycelial nuclei as indicated by co-localization with the mCherry labeled nuclear marker MoHis1-mCherry confirming that they are indeed localized to nuclei as expected. (**A**) MoMyb13-GFP localizes to nuclei. (**B**) MoMyb2-GFP localizes to nuclei but mainly to what is most probably the nucleoli. (**C**) MoMyb15-GFP localizes to nuclei. DIC= Differential Interference Contrast.

### 7. Expression of all identified *MoMyb* genes during infection

All *MoMyb* genes (1-19) were investigated for expression during different HPI, *in vitro* (MY), and in conidia using RT-qPCR. The analysis focuses on the genes’ HPI expression profiles and makes them comparable and was used to make a PCA analysis of correlations of profiles. A minimum spanning tree almost equivalent to clustering is also shown in the resulting PCA plot (**Fig. S5**). The Minimum spanning tree is similar to a more conventional cluster plot using neighbor-joining clustering of the same data (Gower similarity index and final branch as root) (**Fig. 8**). The similarities between profiles can also be seen in a table containing expression profiles for each gene (**Table S2**) and agrees well with blast similarities (**Table S3**). The latter suggests that similar proteins have similar expression profiles.

**Figure 8.**
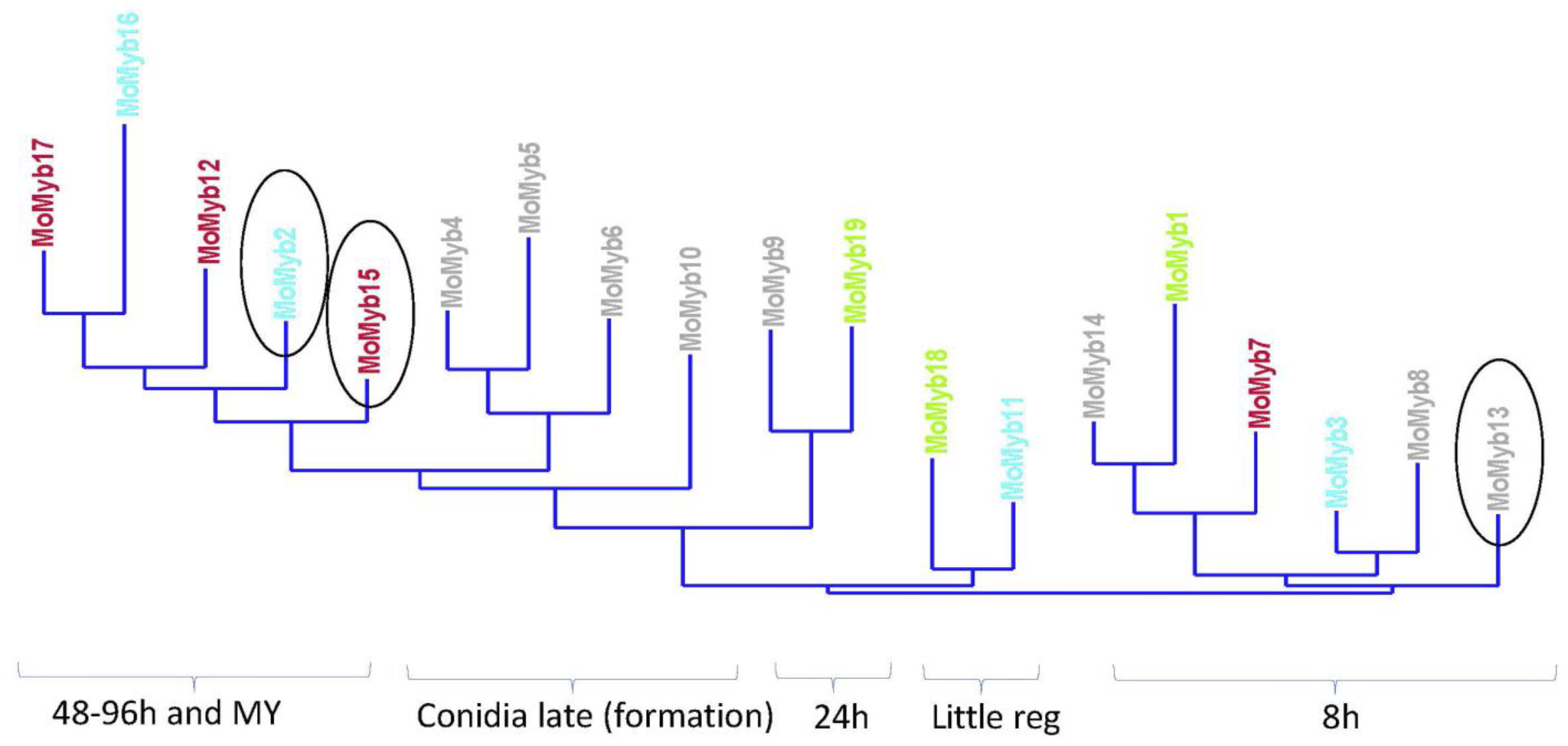
The different *MoMyb* genes show different expression profiles. The expression profiles cluster into 5 groups depending on hours post-infection (HPI) expression during infection. This clustering is, in principle, similar to what can be seen in the PCA plot (**Fig. S5**). Transcripts from growth on agar medium (MY), conidia (late in the formation of conidia), and in rice plant leaves at 8 HPI (8h), 24 HPI (24h), 48 HPI (48h), 72 HPI (72h), and 96 HPI (96h). The color code used is the same as in **Fig. 1B** for the phylogeny clusters showing that the regulatory sub-clusters activated at different HPI contain sets of genes with little similarity in sequence homology. Ellipses mark the 3 genes with phenotypes different from Ku80 for their respective deletion mutants.

MoMyb2 and 15 appear to have essential roles in the necrotrophic stage and in vitro. On the other hand, MoMyb13 appears to be upregulated and used during the early pre-biotrophic stage (8h) of plant contact, indicating that it could be a regulator of penetration and establishment in the plant.

*MoMyb8* is, in our data, upregulated in the early phase and has one peak in the later phases. It seems to be involved in responses to light (Lee et al., 2021). *MoMyb1* has previously been shown to be essential for infection (Dong et al., 2015), and in our data, it belongs to the regulatory cluster upregulated at 8 HPI. Since we could not find growth or infection phenotypes for more than 3 MoMyb proteins encoded by the newly identified genes, it was expected that some of the *MoMyb* genes were not expressed during plant infection at all or pseudogenes, but it turned out that all genes were well expressed and expressed in planta during at different HPIs (**Fig. 8**).

## DISCUSSION

The 19 genes predicted to encode Myb domain-containing proteins in *M. oryzae* (MoMyb1-19) found in the Fungal Transcription Factor Database (FTFD) encodes proteins with very different sizes and content of other domains (**Fig. 1A**). Thus, the Myb domain-containing genes are expected to have diverse functions (Hogues *et al*., 2008; Yu *et al*., 2010; Rodríguez-Sánchez *et al*., 2011; Mukherjee *et al*., 2012; Pattabiraman and Gonda, 2013; Roy, 2016; X. Wang *et al*., 2018; Ma and Constabel, 2019; Ramos-Sáenz *et al*., 2019), and might not all be traditional transcription factors. The latter is often implied for plant Myb proteins (Allan and Espley, 2018; Ambawat et al., 2013; Baldoni et al., 2015; Cao et al., 2020; Dong et al., 2015; Du et al., 2009; Dubos et al., 2010; Kim et al., 2014; Li et al., 2019; Li et al., 2019; Lin et al., 2011; Lin et al., 2012; Liu et al., 2015; Ma and Constabel, 2019; Mateos et al., 2006; Matheis et al., 2017; Mitra, 2018; Pattabiraman and Gonda, 2013; Prouse and Campbell, 2012; Roy, 2016; Valsecchi et al., 2017; Verma et al., 2017; X., Wang et al., 2018; Wieser and Adams, 1995). We found 6 more Myb-domain protein-encoding genes than the 13 *MoMyb* genes identified earlier (Verma et al., 2017) and 9 more than were recently identified (Lee et al., 2021). Interestingly, all 19 MoMyb-protein-encoding genes are expressed at different stages of plant infection (**Fig. 8, S5,** and **Table S2**). Thus, it surprised us that the successful only the deletion of two of our deleted genes (*Δmomyb13* and *ΔmoMyb15*) and one previously deleted gene with a known effect on the infection (*Δmomyb1*) had notable effects on plant infection (**Fig. 3, S3** and **Table1**). There are many possible explanations for this. One explanation is that these deleted genes are not involved in the infection process, as seems to be the case for MoMyb8 (Lee et al., 2021). Nevertheless, since all genes are expressed well in planta at different stages of plant infection (**Table S2**), we suggest that the explanation is redundancy in function or genetic compensation for most of these gene deletions often found negating phenotypic effects of knockouts (El-Brolosy and Stainier, 2017). Similar regulated genes not belonging to the Myb-containing genes or regulated by the Myb genes can create this redundancy or genetic compensation (**Fig. 8, S5,** and **Table S2**). Such genes might collaborate with the deleted genes at different time points or by upregulation of genes with similarity in amino acid sequence with overlapping functions since we found the MoMyb-genes sort into 4 clusters of similarity (**Fig. 1B**). Of course, the 5 genes we could not delete that have not been deleted before (*momyb11, 12, 14, 17, and 19*) could have a role in pathogenicity. If these are genuinely essential for growth in culture, and because of that cannot be deleted, they naturally have a profound effect on pathogenicity since they might be essential for survival also during infection. Interestingly these potentially essential genes are members of different co-regulated clusters identified (**Fig. 8 and S5**), making it possible for them to encode proteins that act together with the other Myb proteins and take over their functions.

**Table 1.** Phenotype summary of MoMybs.

| Protein name | Gene ID | Deletion | Growth | Conidiation | Appressoria formation | Pathogenicity | localization | Stress response | Ref |
| --- | --- | --- | --- | --- | --- | --- | --- | --- | --- |
| MoMyb1 | MGG_06898 | Yes | Reduced | Lost | Reduced | Reduced | cytoplasm and nucleus | Less sensitive to salt and sorbitol | Dong et al., 2015; Lee et al., 2021 |
| MoMyb2 | MGG_10426 | Yes | Reduced | Normal | Normal | Normal | nucleolus | More sensitive to SDS | This study; Lee et al., 2021 |
| MoMyb3 | MGG_08137 | Yes | Normal | Normal | Normal | Normal | - | - | This study; Lee et al., 2021 |
| MoMyb4 | MGG_01426 | Yes | Normal | Normal | Normal | Normal | nucleus | - | This study; Lee et al., 2021 |
| MoMyb5 | MGG_15357 | Yes | Normal | Normal | Normal | Normal | nucleus | - | This study; Lee et al., 2021 |
| MoMyb6 | MGG_05945 | Yes | Normal | Normal | Normal | Normal | - | - | This study; Lee et al., 2021 |
| MoMyb7 | MGG_02746 | Yes | Normal | Normal | Normal | Normal | - | - | Lee et al., 2021 |
| MoMyb8 | MGG_01133 | Yes | Normal | Normal | Normal | Normal | - | - | This study; Lee et al., 2021 |
| MoMyb9 | MGG_05240 | Yes | Normal | Normal | Normal | Normal | nucleus | - | This study; Lee et al., 2021 |
| MoMyb10 | MGG_05748 | Yes | Normal | Normal | Normal | Normal | - | - | This study |
| MoMyb11 | MGG_05099 | Not | - | - | - | - | nucleus | - | This study |
| MoMyb12 | MGG_01130 | Not | - | - | - | - | - | - | This study |
| MoMyb13 | MGG_14558 | Yes | Reduced | Lost | Normal | Lost | nucleus | More sensitive to CR and H<sub>2</sub>O<sub>2</sub>; Less sensitive to NaCl and SOR | This study |
| MoMyb14 | MGG_03899 | Yes | Normal | Normal | Normal | Normal | - | - | This study |
| MoMyb15 | MGG_01012 | Yes | Reduced | Lost | Normal | Lost | nucleus | More sensitive to SDS | This study |
| MoMyb16 | MGG_00138 | Yes | Normal | Normal | Normal | Normal | nucleus | - | This study |
| MoMyb17 | MGG_01720 | Not | - | - | - | - | nucleus | - | This study |
| MoMyb18 | MGG_08095 | Yes | Normal | Normal | Normal | Normal | nucleus | - | This study |
| MoMyb19 | MGG_06120 | Not | - | - | - | - | - | - | This study |

A gene encoding a Myb protein that affected both growth and infection phenotypes is *MoMyb13*.This protein is large (**Fig. 1A**) and has no orthologues outside Ascomycota in the NCBI database. We investigated it further through bioinformatics and found similar large proteins predicted in 19 other published genomes (**Fig. S6**). Three of these hits were in other strains of *Pyricularia oryzae (M. oryzae* teleomorph), which is not strange. Maybe more interesting, common to all hits, including the ones in *M.oryzae*, they are all fungi known to produce melanized hydrophobic structures (Collado et al., 2002; Liu et al., 2009; Kokaew et al., 2011; Al-Khawaldeh et al., 2020; Geisen et al., 2021; Sarsaiya et al., 2020; Li et al., 2016; Gao et al., 2021; Chen et al., 2021) suggesting that a possible involvement of MoMyb13 in hydrophobins formed together with melanin production could be relevant also for these similar proteins in diverse fungi all belonging to the class Sordariomycetes.

Interestingly, all these melanized fungi are known to grow endophytically, as *M. oryzae* do in the first biotrophic stages of infection (Kankanala et al., 2007) when *MoMyb13* is specifically upregulated (**Fig. 8 and Fig. S5).** If these genes and gene products have similar functions as in *M. oryzae* is unknown. Future research should investigate whether these genes from the other fungi could complement *MoMyb13* or each other since *MoMyb13* seems essential in the early biotrophic establishment stage of *M. oryzae* infection of rice. The orthologues of its gene’s products are only present in relatively closely related ascomycetes having biotrophic/endophytic relationships with plants (**Fig. S6**), suggesting that this gene and its product have some essential function for establishing biotrophic interactions. The *MoMyb13* gene or its gene product could also potentially be targeted by more pathogen-specific control measures since it does not seem widely spread among fungi outside the class Sordariomycetes.

The other gene that showed changed strain phenotypes when deleted was *MoMyb15*, which encodes for an ISW2-like protein containing a SANT-type Myb domain. MoMyb15 is at NCBI annotated as an ISW2 type Myb-containing protein conserved in eukaryotes. A true MoISW2 is potentially involved in gene regulation by regulating the access of transcription factors by binding to DNA and controlling nucleosome positioning regulating the access of DNA for other transcription factors and repressors instead of being a transcription factor itself (Fazzio et al., 2005; Hada et al., 2019; Kagalwala et al., 2004). It would be interesting to confirm this function and focus on which DNA sequences it binds to (CHP-seq) and the regulatory effect of the deletion on genes physically adjacent to the DNA binding site in the genome. According to our results (**Fig. 8 and S5 and Table S2**), the *MoMyb15* gene is upregulated at HPIs at the transition from biotrophy to necrotrophy (Kankanala et al., 2007), suggesting it could be a “master regulator” for regulating the fungus many defenses against plant defenses that are strongly upregulated in the biotrophy/necrotrophy transition and during the necrotrophic stage (Kou et al., 2019). Thus, we have initiated detailed research on MoMyb15 to investigate if it has a “master regulator” function working through nucleosome condensation in *M. oryzae*.

The genes encoding proteins with Myb-domains showing no phenotypes when deleted should also be subject to studies since they were upregulated at specific infection stages. However, these genes could be studied using knockdown systems to avoid mutation-induced genetic compensation (El-Brolosy and Stainier, 2017). Conditional knockdown systems can maybe be used for essential genes we did not manage to knock out, even if the most common TET-on TET-off can create artifacts due to mitochondrial poisoning (Moullan et al., 2015).

## METHODS

### Organisms and media used

*Magnaporthe oryzae* B. Couch anamorph of the teleomorph *Pyricularia oryzae* Cavara was used for this research. As background *M. oryzae* strain, we used Ku80 to minimize random integration events when transformed (Villalba et al., 2008). The susceptible Indica rice (cv. CO-39) and barley (cv. Golden Promise) used for the fungal pathogenicity tests were from the seed bank of our laboratory. The CM (complete medium), MM (minimal medium), and RBM (rice bran medium) used for growing the fungus were prepared as described (Li et al., 2019). The *Escherichia coli* strain DH-5α used for routine bacterial transformations (Li et al., 2015) and maintenance of various plasmid vectors was bought from Solarbio Life Sciences, China.

### Knockouts, complementations, and verifications

The *Myb* gene deletion vectors were constructed by inserting the 1 kb up-and-down-stream fragments of the target gene’s coding region as the flanking regions of the HPH (hygromycin phosphotransferase) gene of the plasmid pBS-HYG (Li et al., 2012). No less than 2 μg of the deletion vector DNA of the target gene was introduced to Ku80 protoplasts, and transformants were selected for hygromycin resistance to perform gene deletion transformations. Southern blotting was conducted to confirm the correct deletion using the digoxigenin (DIG) high prime DNA labeling and detection starter Kit I (11745832910 Roche Germany). The *Myb* gene complementation vectors were constructed by cloning the entire length of the target gene with the native promoter region (about 1.5-kb) to the pCB1532 plasmid. When making the complementation vector, *GFP* was linked to the C-terminal of the target genes to study the sub-cellar localization of Myb proteins. The constructed vector DNA was introduced into the mutation protoplast for the gene complementation assay, and the transformants were screened using 50 μg/ml chlorimuron-ethyl to select successful complementation strains. The detailed fungal protoplast preparation and transformation methods have been described previously (Li et al., 2016). All primers needed for these knockouts and complementations are listed (**Table S1**). The sub-cellar localization of Myb proteins was observed by confocal microscopy (Nikon A1). GFP and RFP excitation wavelengths were 488 nm and 561 nm, respectively.

### Colony and growth phenotypes measurements

Vegetative growth was tested by measuring the colony diameter after ten days of growth in 9 cm Petri dishes at 25°C under 12h-to-12h light and dark periods. Conidia production was evaluated by flooding the 12-day-old colony with double distilled water, filtering out the mycelia with gauze, and counting the conidia using a hemacytometer. The conidiophore induction assay was performed by excising one thin agar block from the fungal colony and then incubating it in a sealed chamber for 24 h with constant light (Li et al., 2010b). Mycelia appressoria were induced by placing a suspension of mycelial fragments on a hydrophobic surface in a humid environment at 25°C for 24h. The pathogenicity assay on rice was performed by spraying 5 ml conidial suspension (5 ×104 spores/ml) on 15-day-old plants (Li et al., 2014). Post-spraying inoculated plants were kept in a sealed chamber with a 90% relative humidity at 25°C for 24 h. Then, the inoculated plants were removed from the chamber to allow disease symptoms to develop for 4-5 days. The pathogenicity assay on excised barley and rice leaves was performed by cutting a small block from the agar culture of the fungus and placing it on excised leaves for five days in a moist chamber for disease development (Li et al., 2010a). The sexual reproduction was performed by crossing the tested strain with the sexually compatible strain TH3 in OM plates and then incubating at 19°C for 30 days with continuous light. The perithecia and clavate asci were photographed by a microscope equipped with a camera (OLYMPUS BX51).

### RT-qPCR

Total RNA was extracted using Eastep®Super Total RNA Extraction Kit (Promega (Beijing) Biotechnology, LS1040) to perform RT-qPCR, and 5 mg of RNA was reverse-transcribed to cDNA using the Evo M-MLV RT kit with gDNA to clean before the qPCR (Accurate Biotechnology (Hunan), AG11705) according to the manufacturer’s instructions. The resulting cDNA was diluted ten times and used as the template for qPCR. The qPCR reactions were performed using an Applied Biosystems 7500 Real-Time PCR System. Each reaction contained 25 ul of SuperRealPreMix Plus SYBR Green (Tiangen Biotechnology, Beijing, FP205-02), one μl of cDNA, and 1.5 μl of each primer solution. The thermal cycling conditions were 15 min at 95 °C followed by 40 cycles of 10 s at 95 °C and 20 s at 60 °C. The threshold cycle (Ct) values were obtained by analyzing amplification curves with a normalized reporter threshold of 0.1. The relative expression value was calculated using the 2^-ΔΔCT^ method (Livak and Schmittgen, 2001).

### PCA analysis and clustering of RT-qPCR expression data

First, expression values were normalized using the beta-tubulin gene as the housekeeping reference gene. Since we did not want to give too much weight to the very artificial conditions in lab media for mycelium growth (MY), average expression at all the measured HPIs was used for normalization to focus on the *in planta* variation at different HPIs and to make comparisons of gene expression profiles during infection of rice more critical for the analysis. For the PCA, we used correlation of the data for the different genes since the genes have different averages of expression. Together that focuses the analysis on the genes’ varying HPI profiles and makes them comparable. The analysis is shown as a PCA bi-plot, of the PC1 and PC2 with gene names as objects, the variables vectorial contributions to the PCs (the different time HPIs, MY=mycelium in culture, and Con = conidia before inoculation) (**Fig. S5).** We used the freeware PAST (Hammer et al., 2001) version 4.08 (released November 2021) for the PCA analysis. It is available from the University of Oslo, Natural History Museum https://www.nhm.uio.no/english/research/infrastructure/past/. The data was entered and handled in MS Excel and then copy-pasted into PAST for analysis. The same data is, in addition, presented using the clustering function in PAST to present the data in a more usual way using a clustering method that gives a similar result as the Minimum spanning tree (Neighbor-joining, Gower similarity, Final branch as root) (**Fig. 8**) and presented with the same color coding as in **Figure 1b** for comparison.

## Author contributions

YL, domain prediction and evolutionary analysis, Myb gene deletions, phenotype tests, protein localization, RT-qPCR, data collection and analysis, figure making, manuscript preparation, and writing. XZ, MoMyb13 gene deletion, MoMyb13 mutant phenotype analysis, MoMyb13 protein localization, RT-qPCR of hydrophobin gene. MC, MP, CL, LG, PH, Myb-gene deletions. SZ, RT-qPCR of all Myb genes, protein localization of 8 Myb genes. SO, RT-qPCR extended analysis including PCA and clustering, extended analysis of possible roles of Myb13 and Myb15, manuscript preparation, and writing.

## Acknowledgments

The strain Ku80 used in this study was obtained initially from Nicholas J. Talbot, University of Exeter, UK. This research was financially supported by the National Science Foundation of China (NSFC31871914)

## DATA AVAILABILITY STATEMENT

Data supporting the findings of this study are available within the paper and its supplements.

## Figure Legends

**Figure S1.**
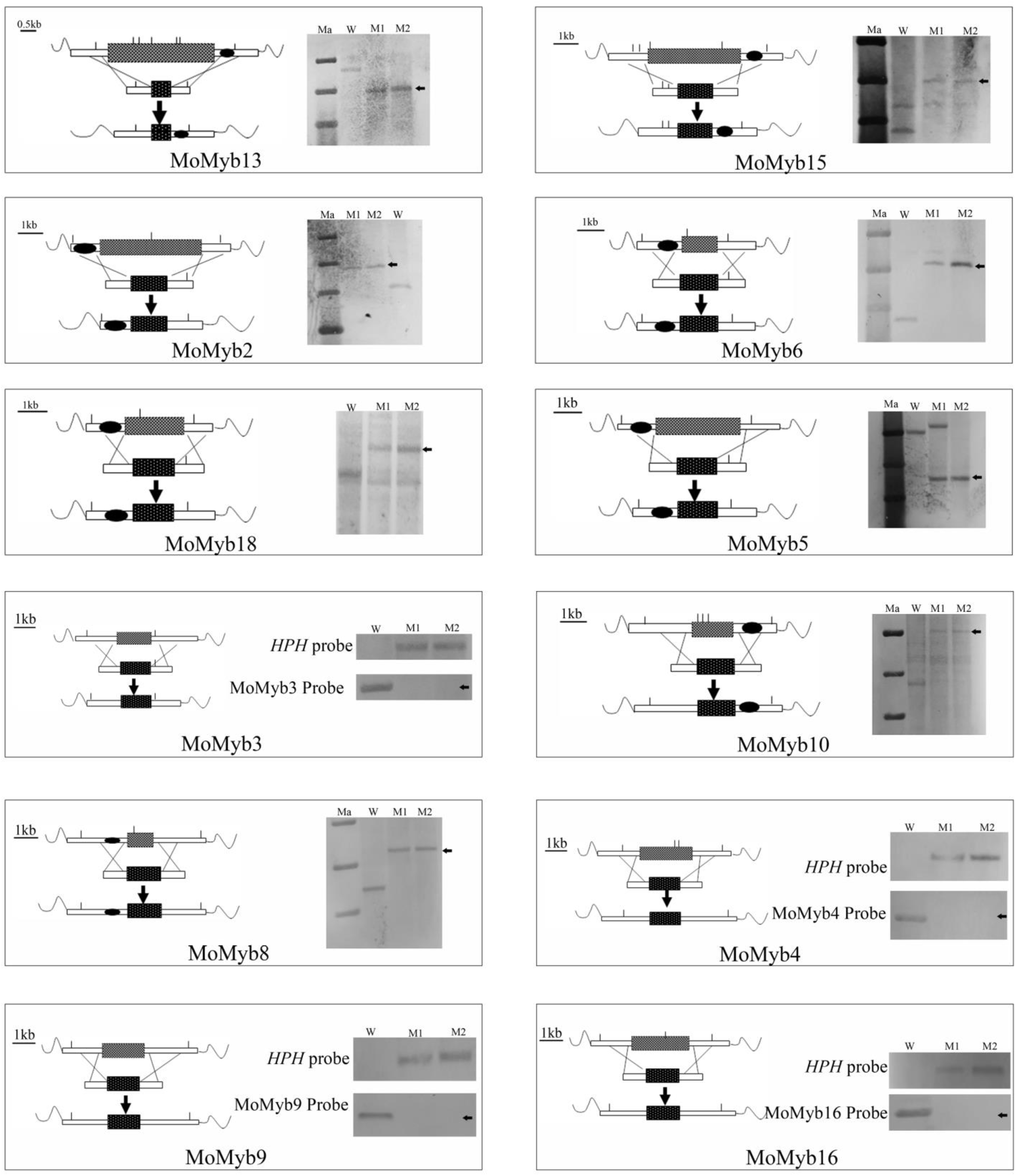
Southern blot confirmation for all the 12 mutants obtained in this study. The HPH gene or the 1.0 kb up- or downstream of the target gene was used as a probe to detect successful gene knockout events.

**Figure S2.**
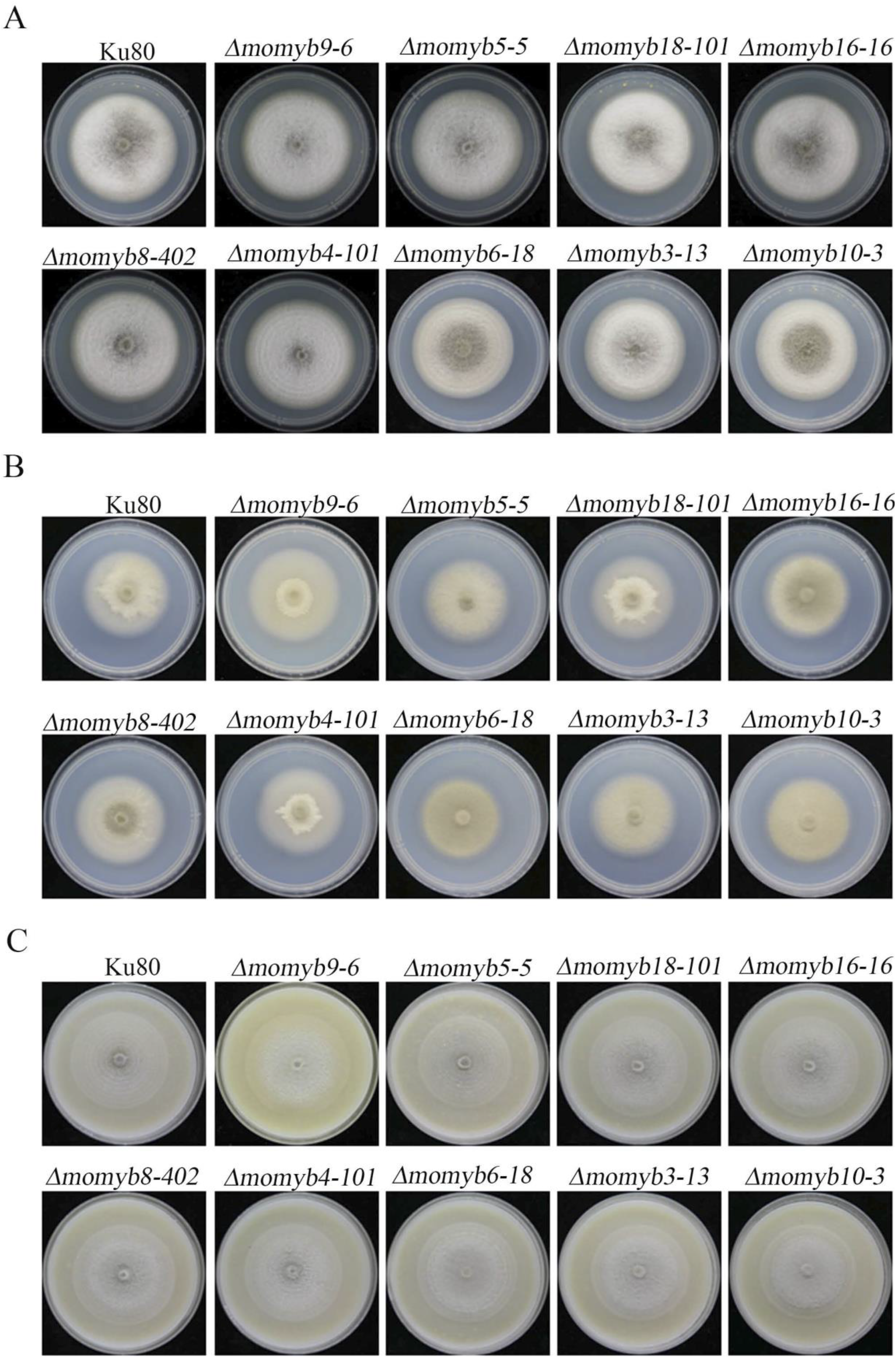
Colony morphology of the 9 *Myb* gene mutants with no effect on pathogenicity compared to Ku80 on CM (A) MM (B), and RBM (C).

**Figure S3.**
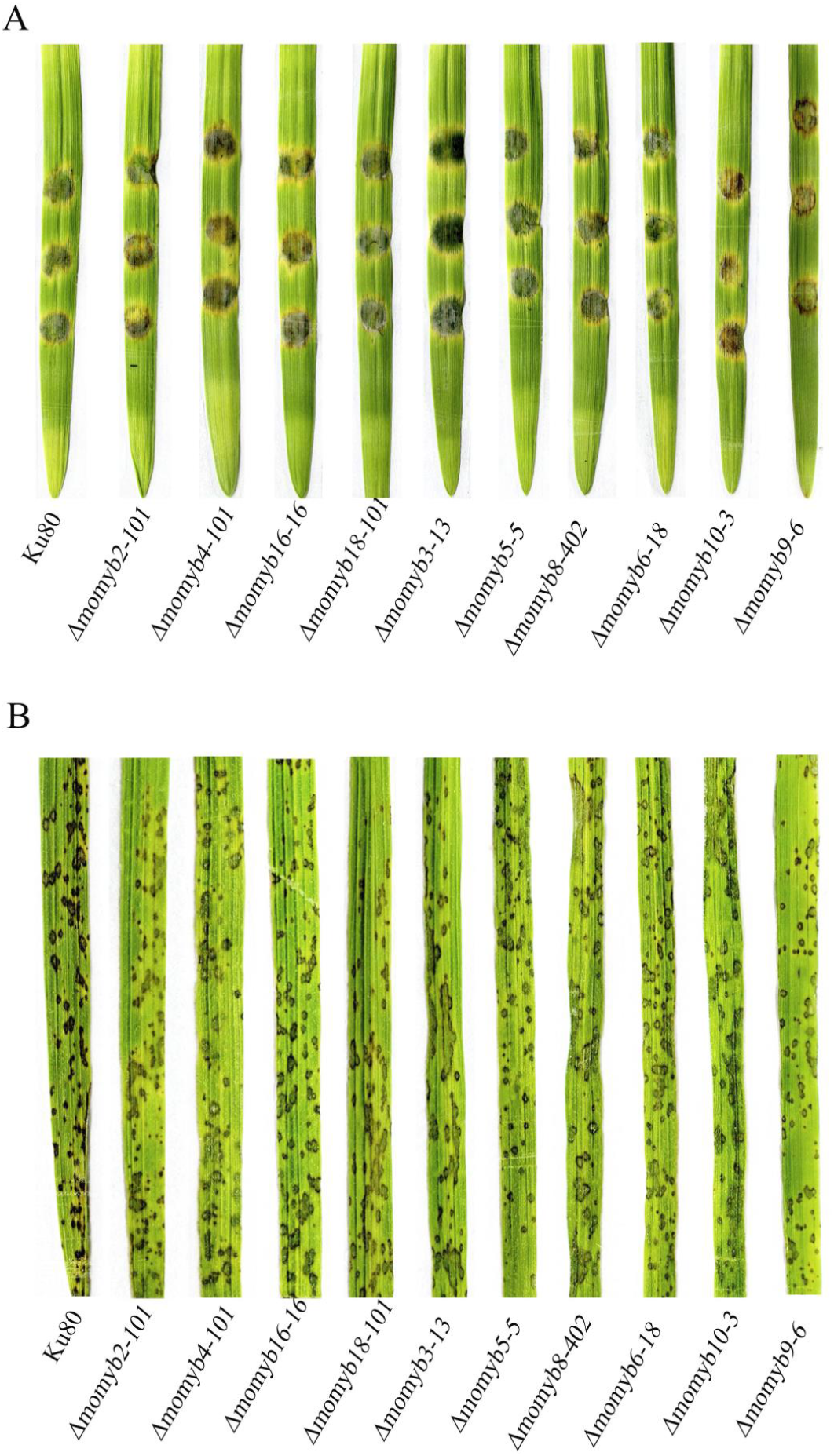
No effect on pathogenicity due to the *Myb* gene mutations can be seen for 9 mutants tested using either the agar block technique or conidia sprayed on the leaves. (**A**) Test of mutants and background strain (Ku80) on barley leaves using the agar block technique (**B**) Test of mutants and background strain (Ku80) conidia sprayed on rice leaves.

**Figure S4.**
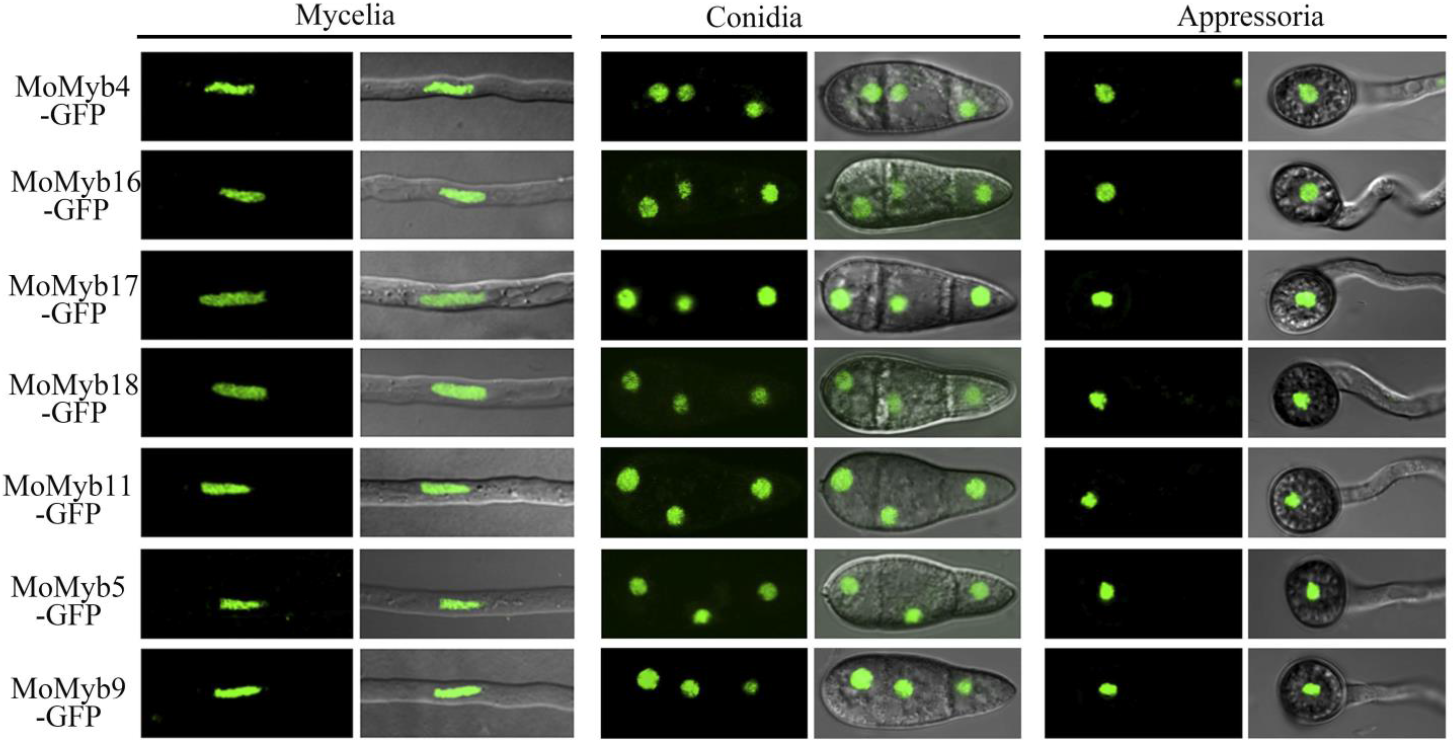
Localization of the MoMyb-GFP complements from which we could find strong enough GFP signals in mycelia conidia and appressoria. The localization pattern suggests nuclear localisations as expected for Myb-domain-containing proteins.

**Figure S5.**
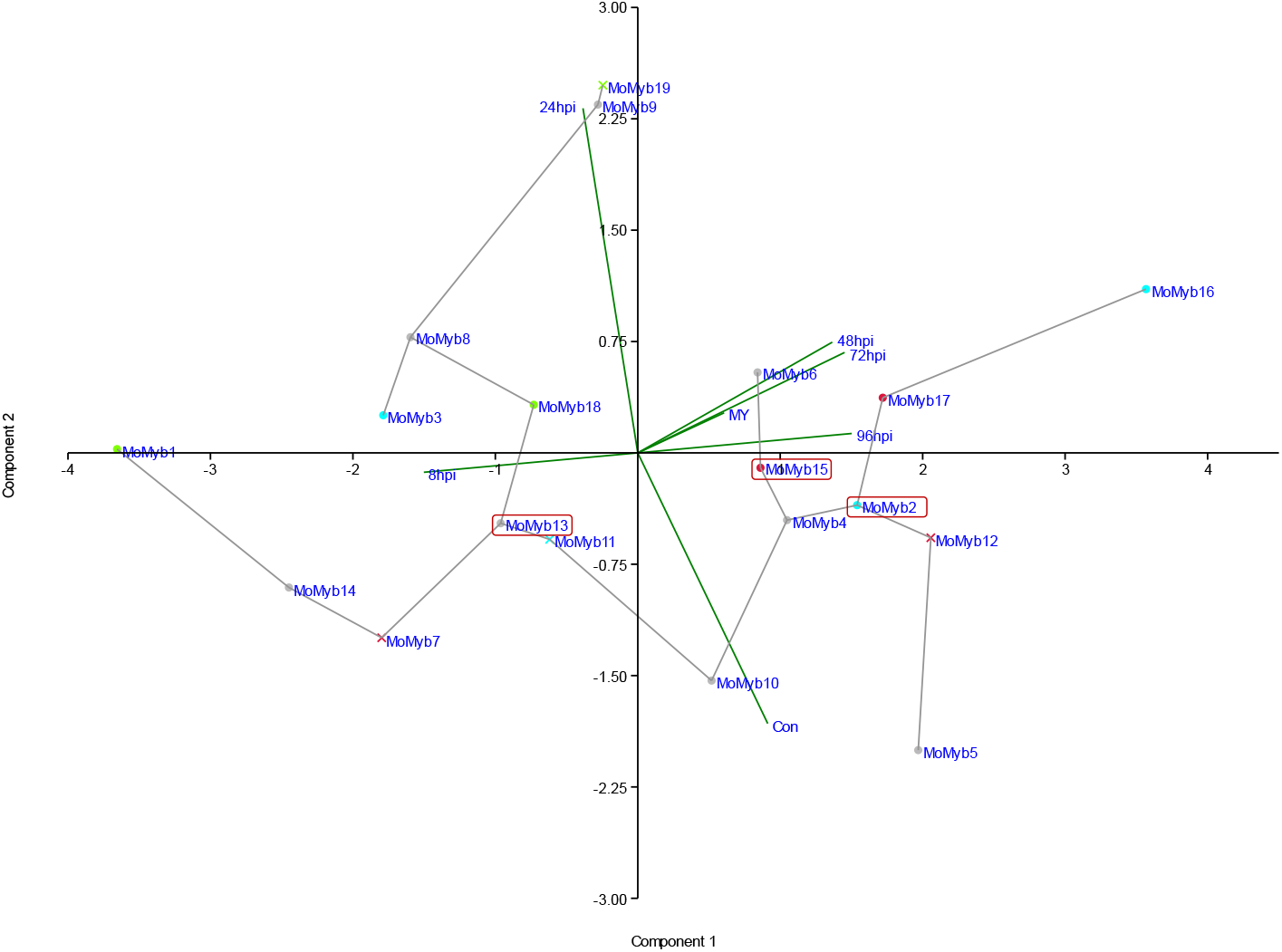
The *MoMyb* genes show different expression profiles visualized by Principal Component Analysis (PCA). Variables were transcription with different treatments and HPIs from infected rice leaves, growth on agar medium (MY), conidia (Conidia late), 8 HPI (8h), 24 HPI (24h), 48 HPI (48h), 72 HPI (72h) and 96 HPI (96h). The expression values of the 19 identified genes encoding Myb domain-containing proteins were the objects. The first and second PCA are presented after a PCA based on the correlation of the expression profiles (as shown in **Table S2**). The plot is a biplot also showing the loading of the variables in the two PCs. A “Minimum spanning tree” nearly identical to the tree in Fig. 8 joins the most similar expressed genes for all calculated PCs. The color code used for the marker dots (or Xs) in front of the gene names is the same as in **Fig. 1B** for the predicted amino acid sequence similarity clusters, showing that the regulatory clusters contain sets of genes with little similarity in sequence homology. A marker X in front of the gene name shows a gene we could not delete. Boxes mark the 3 genes with noted phenotypes for their deletion mutants. MoMyb1, furthest to the left in the figure, is known from the literature to be activated early in infection, and here it is far left in component 1 with the 8 HPI negative loading corroborating the previous results from the literature.

**Figure S6.**
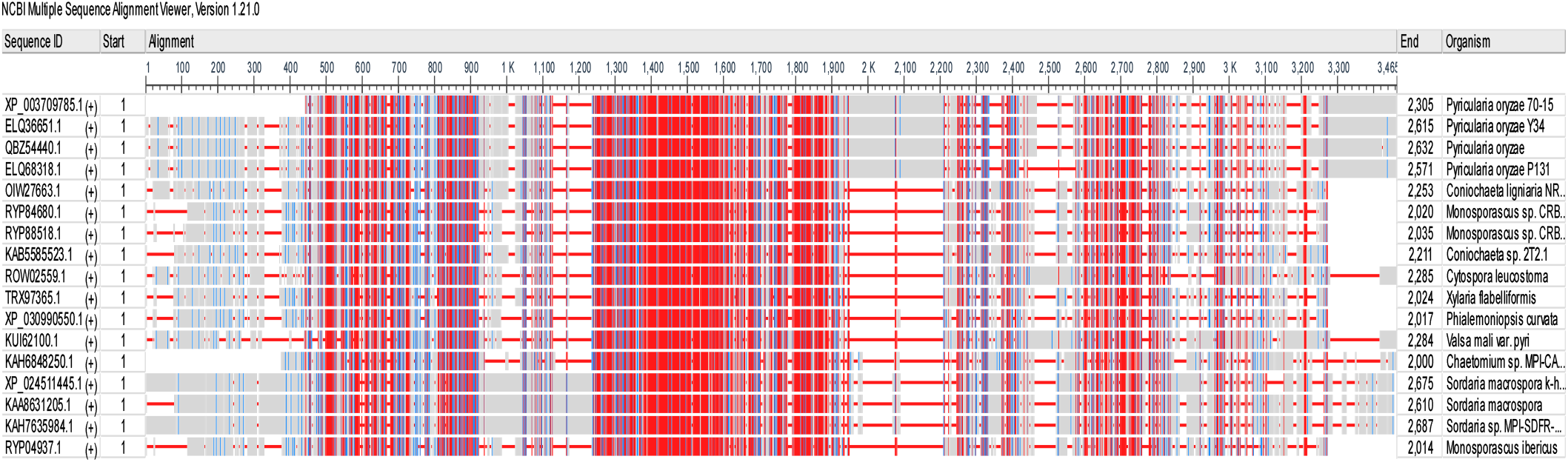
Alignments of the most similar orthologues to the MoMyb13 protein from species related to *M. oryzae (Pyricularia oryzae*)(all are with E-values less or equal to 3E-135 and with more than 70% query cover) (**Table S3**). Data from an NCBI produced by the alignment of all fungal genomes annotated.

**Table S1.** Primers used in this study, their sequences, and what they were used to accomplish.

| Protein name | Gene ID | Deletion | Growth | Conidiation | Appressoria formation | Pathogenicity | localization | Stress response | Ref |
| --- | --- | --- | --- | --- | --- | --- | --- | --- | --- |
| MoMyb1 | MGG_06898 | Yes | **Reduced** | **Lost** | **Reduced** | **Reduced** | cytoplasm and nucleus | Less sensitive to salt and sorbitol | Dong et al., 2015; Lee et al., 2021 |
| MoMyb2 | MGG_10426 | Yes | **Reduced** | Normal | Normal | Normal | nucleolus | More sensitive to SDS | This study; Lee et al., 2021 |
| MoMyb3 | MGG_08137 | Yes | Normal | Normal | Normal | Normal | - | - | This study; Lee et al., 2021 |
| MoMyb4 | MGG_01426 | Yes | Normal | Normal | Normal | Normal | nucleus | - | This study; Lee et al., 2021 |
| MoMyb5 | MGG_15357 | Yes | Normal | Normal | Normal | Normal | nucleus | - | This study; Lee et al., 2021 |
| MoMyb6 | MGG_05945 | Yes | Normal | Normal | Normal | Normal | - | - | This study; Lee et al., 2021 |
| MoMyb7 | MGG_02746 | Yes | Normal | Normal | Normal | Normal | - | - | Lee et al., 2021 |
| MoMyb8 | MGG_01133 | Yes | Normal | Normal | Normal | Normal | - | - | This study; Lee et al., 2021 |
| MoMyb9 | MGG_05240 | Yes | Normal | Normal | Normal | Normal | nucleus | - | This study; Lee et al., 2021 |
| MoMyb10 | MGG_05748 | Yes | Normal | Normal | Normal | Normal | - | - | This study |
| MoMyb11 | MGG_05099 | Not | - | - | - | - | nucleus | - | This study |
| MoMyb12 | MGG_01130 | Not | - | - | - | - | - | - | This study |
| MoMyb13 | MGG_14558 | Yes | **Reduced** | **Lost** | Normal | **Lost** | nucleus | More sensitive to CR and H<sub>2</sub>O<sub>2</sub>; Less sensitive to NaCl and SOR | This study |
| MoMyb14 | MGG_03899 | Yes | Normal | Normal | Normal | Normal | - | - | This study |
| MoMyb15 | MGG_01012 | Yes | **Reduced** | **Lost** | **Normal** | **Lost** | nucleus | More sensitive to SDS | This study |
| MoMyb16 | MGG_00138 | Yes | Normal | Normal | Normal | Normal | nucleus | - | This study |
| MoMyb17 | MGG_01720 | Not | - | - | - | - | nucleus | - | This study |
| MoMyb18 | MGG_08095 | Yes | Normal | Normal | Normal | Normal | nucleus | - | This study |
| MoMyb19 | MGG_06120 | Not | - | - | - | - | - | - | This study |

**Table S1.**
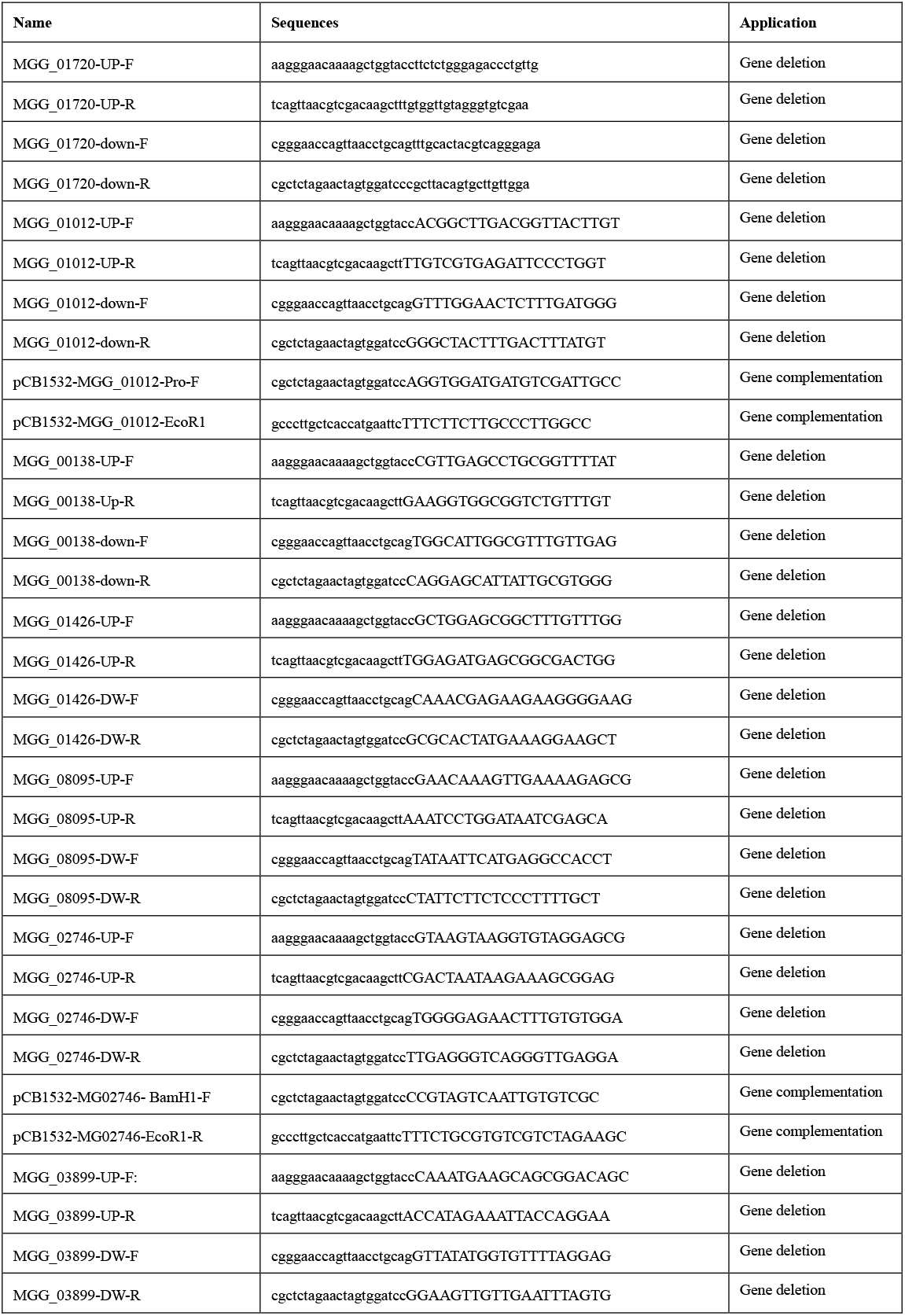
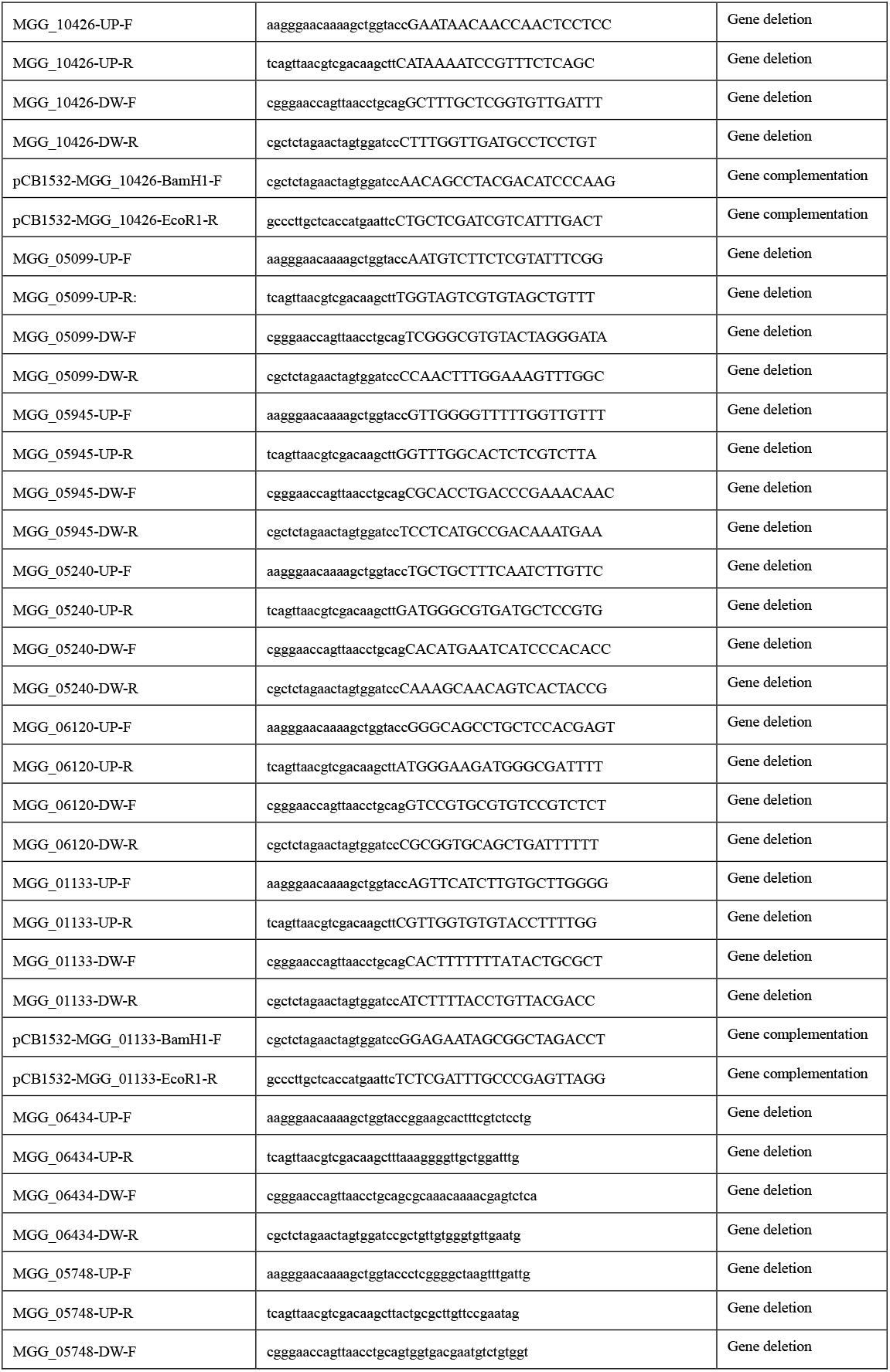
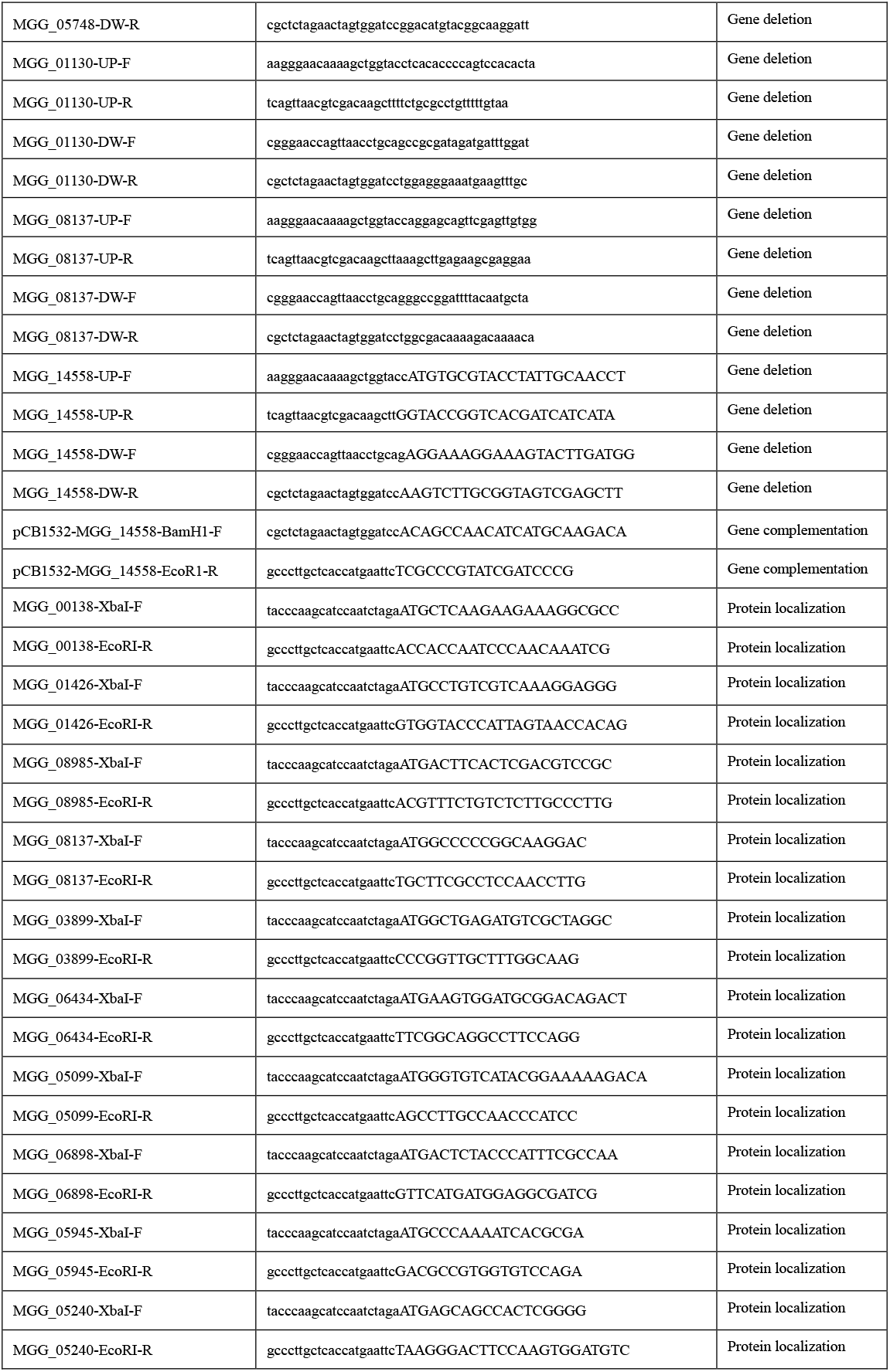
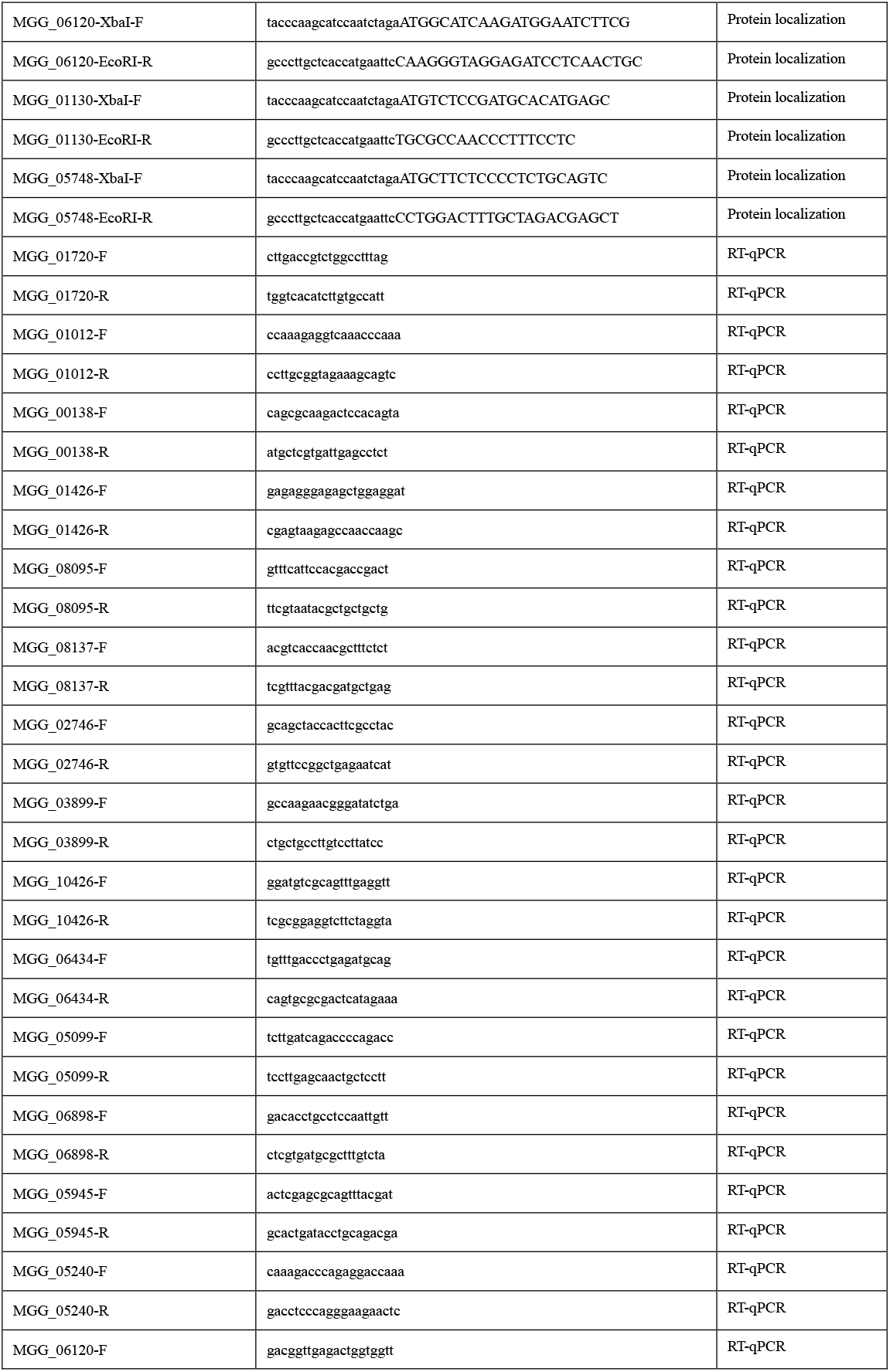
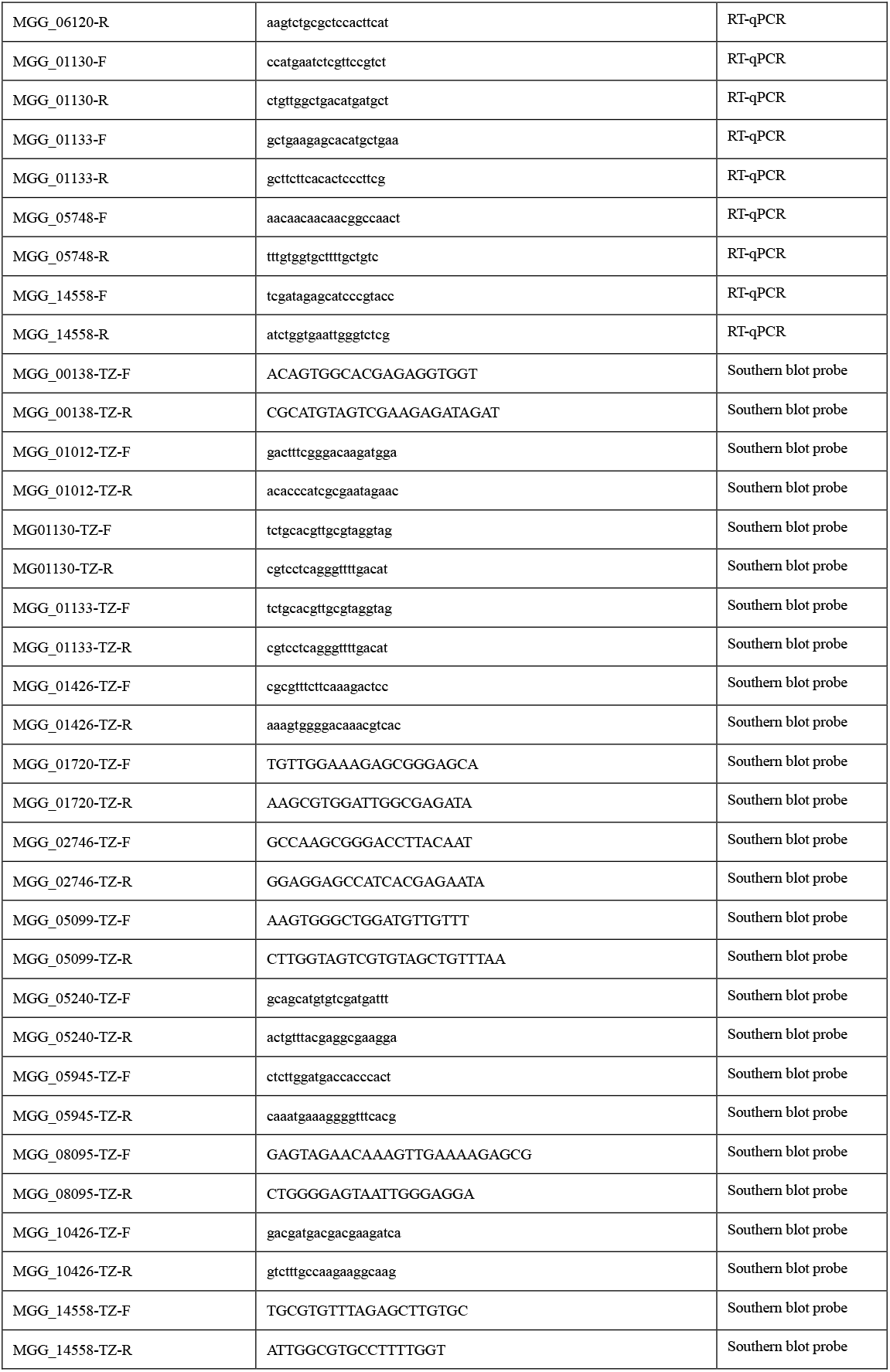
Primers used in this study.

**Table S2.**
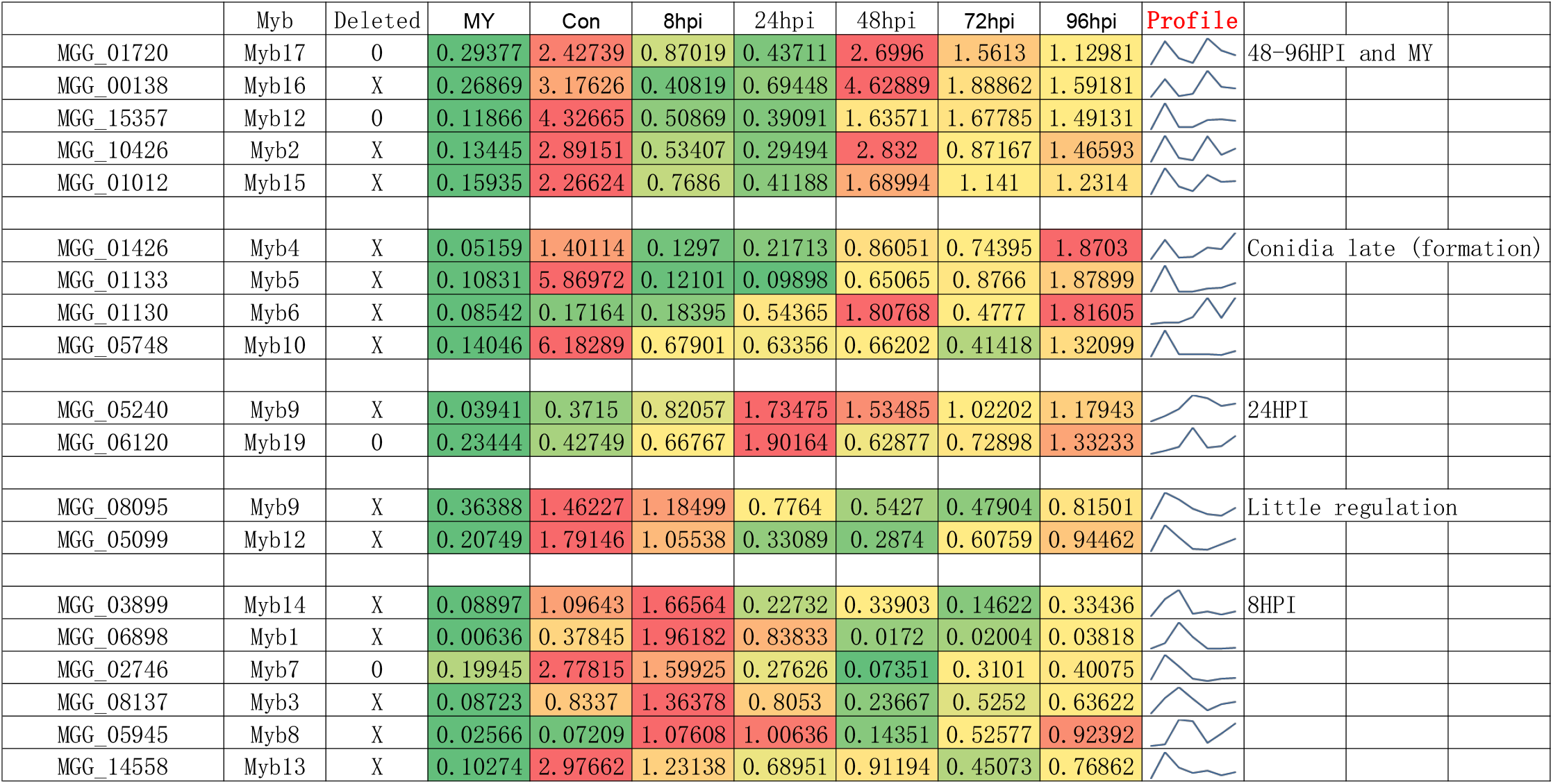
List of the MoMyb proteins organized in groups after their judged expression profiles. Deleted genes in this study are marked by an X in or a 0 if not deleted. The relative expression in normalized by the average expression during infection (HPI8-96) for each gene (the HPI columns) shown in columns. MY=in vitro mycelium), Con= conidia, then 8-96HPI columns. Profile shows the profile My-Con-8hpi-24hpi-48hpi-72hpi-96hpi. The transition from biotrophy to necrotrophy occurs at around 48hpi or just before. Infection and entry into biotrophy are during the first 24 hours. The two first, MY and Con shows expression in vitro and conidia before penetration of the leaf. Comments to the far right tell when the respective groups of genes are relatively highly expressed compared to their respective averages during infection.

|  | Myb | Deleted | MY | Con | 8hpi | 24hpi | 48hpi | 72hpi | 96hpi | Profile |  |
| --- | --- | --- | --- | --- | --- | --- | --- | --- | --- | --- | --- |
| MGG_01720 | Myb17 | 0 | 0.29377 | 2.42739 | 0.87019 | 0.43711 | 2.6996 | 1.5613 | 1.12981 |  | 48–96HPI and MY |
| MGG_00138 | Myb16 | X | 0.26869 | 3.17626 | 0.40819 | 0.69448 | 4.62889 | 1.88862 | 1.59181 |  |  |
| MGG_15357 | Myb12 | 0 | 0.11866 | 4.32665 | 0.50869 | 0.39091 | 1.63571 | 1.67785 | 1.49131 |  |  |
| MGG_10426 | Myb2 | X | 0.13445 | 2.89151 | 0.53407 | 0.29494 | 2.832 | 0.87167 | 1.46593 |  |  |
| MGG_01012 | Myb15 | X | 0.15935 | 2.26624 | 0.7686 | 0.41188 | 1.68994 | 1.141 | 1.2314 |  |  |
| MGG_01426 | Myb4 | X | 0.05159 | 1.40114 | 0.1297 | 0.21713 | 0.86051 | 0.74395 | 1.8703 |  | Conidia late (formation) |
| MGG_01133 | Myb5 | X | 0.10831 | 5.86972 | 0.12101 | 0.09898 | 0.65065 | 0.8766 | 1.87899 |  |  |
| MGG_01130 | Myb6 | X | 0.08542 | 0.17164 | 0.18395 | 0.54365 | 1.80768 | 0.4777 | 1.81605 |  |  |
| MGG_05748 | Myb10 | X | 0.14046 | 6.18289 | 0.67901 | 0.63356 | 0.66202 | 0.41418 | 1.32099 |  |  |
| MGG_05240 | Myb9 | X | 0.03941 | 0.3715 | 0.82057 | 1.73475 | 1.53485 | 1.02202 | 1.17943 |  | 24HPI |
| MGG_06120 | Myb19 | 0 | 0.23444 | 0.42749 | 0.66767 | 1.90164 | 0.62877 | 0.72898 | 1.33233 |  |  |
| MGG_08095 | Myb9 | X | 0.36388 | 1.46227 | 1.18499 | 0.7764 | 0.5427 | 0.47904 | 0.81501 |  | Little regulation |
| MGG_05099 | Myb12 | X | 0.20749 | 1.79146 | 1.05538 | 0.33089 | 0.2874 | 0.60759 | 0.94462 |  |  |
| MGG_03899 | Myb14 | X | 0.08897 | 1.09643 | 1.66564 | 0.22732 | 0.33903 | 0.14622 | 0.33436 |  | 8HPI |
| MGG_06898 | Myb1 | X | 0.00636 | 0.37845 | 1.96182 | 0.83833 | 0.0172 | 0.02004 | 0.03818 |  |  |
| MGG_02746 | Myb7 | 0 | 0.19945 | 2.77815 | 1.59925 | 0.27626 | 0.07351 | 0.3101 | 0.40075 |  |  |
| MGG_08137 | Myb3 | X | 0.08723 | 0.8337 | 1.36378 | 0.8053 | 0.23667 | 0.5252 | 0.63622 |  |  |
| MGG_05945 | Myb8 | X | 0.02566 | 0.07209 | 1.07608 | 1.00636 | 0.14351 | 0.52577 | 0.92392 |  |  |
| MGG_14558 | Myb13 | X | 0.10274 | 2.97662 | 1.23138 | 0.68951 | 0.91194 | 0.45073 | 0.76862 |  |  |

**Table S3.** Sequences producing significant alignments with more than 70 percent cover: The similarity of NCBI hit data of all highly similar orthologues to MoMyb13 (E value less or equal 3E-135), other strains of M. oryzae (Pyricularia oryzae) excluded. With more than 70% query cover.

Sequences producing significant alignments with more than 70 percent cover:
| Sequences | Description | Scientific Name | Max Score | Total Score | Query Cover | E value | Per. Ident | Acc. Len | Accession |
| --- | --- | --- | --- | --- | --- | --- | --- | --- | --- |
| Select seq<br>ref\|XP_003709785.1\| | uncharacterized protein MGG_14558 [Pyricularia oryzae 70-15] | Pyricularia oryzae 70-15 | 4526 | 4526 | 100% | 0 | 100.00% | 2305 | XP_003709785.1 |
| Select seq<br>gb\|ELQ36651.1\| | hypothetical protein OOU_Y34scaffold00649g34 [Pyricularia oryzae Y34] | Pyricularia oryzae Y34 | 4510 | 4510 | 100% | 0 | 99.91% | 2615 | ELQ36651.1 |
| Select seq<br>gb\|QBZ54440.1\| | hypothetical protein PoMZ_10140 [Pyricularia oryzae] | Pyricularia oryzae | 4444 | 4444 | 100% | 0 | 98.11% | 2632 | QBZ54440.1 |
| Select seq<br>gb\|ELQ68318.1\| | hypothetical protein OOW_P131scaffold00255g20 [Pyricularia oryzae P131] | Pyricularia oryzae P131 | 4404 | 4404 | 100% | 0 | 97.87% | 2571 | ELQ68318.1 |
| Select seq<br>gb\|ROW02559.1\| | hypothetical protein VPNG_07863 [Cytospora leucostoma] | Cytospora leucostoma | 555 | 681 | 90% | 7.00E-158 | 37.34% | 2285 | ROW02559.1 |
| Select seq<br>gb\|KUI62100.1\| | hypothetical protein VP1G_09221 [Valsa mali var. pyri] | Valsa mali var. pyri | 530 | 659 | 86% | 2.00E-149 | 36.63% | 2284 | KUI62100.1 |
| Select seq<br>gb\|OIW27663.1\| | hypothetical protein CONLIGDRAFT_682689 [Coniochaeta ligniaria NRRL 30616] | Coniochaeta ligniaria NRRL 30616 | 614 | 723 | 82% | 5.00E-178 | 39.22% | 2253 | OIW27663.1 |
| Select seq<br>gb\|KAB5585523.1\| | hypothetical protein GE09DRAFT_29357 [Coniochaeta sp. 2T2.1] | Coniochaeta sp. 2T2.1 | 579 | 690 | 78% | 4.00E-166 | 38.69% | 2211 | KAB5585523.1 |
| Select seq<br>gb\|KAA8631205.1\| | hypothetical protein SMACR_04153 [Sordaria macrospora] | Sordaria macrospora | 506 | 782 | 78% | 3.00E-141 | 43.08% | 2610 | KAA8631205.1 |
| Select seq<br>gb\|KAH7635984.1\| | hypothetical protein B0T09DRAFT_31245 [Sordaria sp. MPI-SDFR-AT-0083] | Sordaria sp. MPI-SDFR-AT-0083 | 506 | 781 | 78% | 5.00E-141 | 42.98% | 2687 | KAH7635984.1 |
| Select seq<br>gb\|TRX97365.1\| | hypothetical protein FHL15_001643 [Xylaria flabelliformis] | Xylaria flabelliformis | 535 | 609 | 77% | 4.00E-152 | 36.34% | 2024 | TRX97365.1 |
| Select seq<br>gb\|RYP84680.1\| | hypothetical protein DL769_001114 [Monosporascus sp. CRB-8-3] | Monosporascus sp. CRB-8-3 | 604 | 664 | 76% | 7.00E-176 | 37.46% | 2020 | RYP84680.1 |
| Select seq<br>gb\|RYP88518.1\| | hypothetical protein DL770_004599 [Monosporascus sp. CRB-9-2] | Monosporascus sp. CRB-9-2 | 598 | 652 | 76% | 2.00E-173 | 36.53% | 2035 | RYP88518.1 |
| Select seq<br>ref\|XP_030990550.1\| | uncharacterized protein E0L32_009657 [Phialemoniopsis curvata] | Phialemoniopsis curvata | 531 | 934 | 75% | 5.00E-151 | 42.68% | 2017 | XP_030990550.1 |
| Select seq<br>ref\|XP_024511445.1\| | uncharacterized protein SMAC_04153 [Sordaria macrospora k-hell] | Sordaria macrospora k-hell | 508 | 780 | 73% | 1.00E-141 | 43.08% | 2675 | XP_024511445.1 |
| Select seq<br>gb\|KAH6848250.1\| | hypothetical protein B0I37DRAFT_159719 [Chaetomium sp. MPI-CAGE-AT-0009] | Chaetomium sp. MPI-CAGE-AT-0009 | 509 | 768 | 72% | 2.00E-143 | 40.27% | 2000 | KAH6848250.1 |
| Select seq<br>gb\|RYP04937.1\| | hypothetical protein DL764_004149 [Monosporascus ibericus] | Monosporascus ibericus | 484 | 692 | 71% | 3.00E-135 | 40.63% | 2014 | RYP04937.1 |

